# Unique and distinct identities and functions of leaf phloem cells revealed by single cell transcriptomics

**DOI:** 10.1101/2020.09.11.292110

**Authors:** Ji-Yun Kim, Efthymia Symeonidi, Tin Yau Pang, Tom Denyer, Diana Weidauer, Margaret Bezrutczyk, Manuel Miras, Nora Zöllner, Michael M. Wudick, Martin Lercher, Li-Qing Chen, Marja C.P Timmermans, Wolf B. Frommer

**Affiliations:** Institute for Molecular Physiology and Cluster of Excellence on Plant Sciences (CEPLAS), Heinrich-Heine-University Düsseldorf, Düsseldorf 40225, Germany; Center for Plant Molecular Biology, University of Tübingen, Auf der Morgenstelle 32, 72076 Tübingen, Germany; Institute for Computer Science and Department of Biology, Heinrich-Heine-University Düsseldorf, Düsseldorf 40225, Germany; Institute of Transformative Bio-Molecules (WPI-ITbM), Nagoya University, Chikusa, Nagoya 464-8601, Japan; Department of Plant Biology, School of Integrative Biology, University of Illinois at Urbana-Champaign, Urbana, Illinois 61801, U.S.A

**Author notes:** Correspondence and requests for materials should be addressed to Ji-Yun Kim.

**Keywords:** Arabidopsis, phloem loading, SWEET transporters, sugar transport, amino acid transport, amino acid metabolism, phloem parenchyma, companion cells

## Abstract

The leaf vasculature plays a key role in solute translocation. Veins consist of at least seven distinct cell types, with specific roles in transport, metabolism, and signaling. Little is known about the vascular cells in leaves, in particular the phloem parenchyma (PP). PP effluxes sucrose into the apoplasm as a basis for phloem loading; yet PP has only been characterized microscopically. Here, we enriched vascular cells from Arabidopsis leaves to generate a single-cell transcriptome atlas of leaf vasculature. We identified ≥19 cell clusters, encompassing epidermis, guard cells, hydathodes, mesophyll, and all vascular cell types, and used metabolic pathway analysis to define their roles. Clusters comprising PP cells were enriched for transporters, including *SWEET11* and *SWEET12* sucrose and UmamiT amino acid efflux carriers. PP development occurs independently from APL, a transcription factor required for phloem differentiation. PP cells have a unique pattern of amino acid metabolism activity distinct from companion cells (CC), explaining differential distribution/metabolism of amino acids in veins. The kinship relation of the vascular clusters is strikingly similar to the vein morphology, except for a clear separation of CC from the other vascular cells including PP. In summary, our scRNA-seq analysis provides a wide range of information into the leaf vasculature and the role and relationship of the leaf cell types.

## Introduction

A key feature of multicellularity is the division of labor. During evolution, when plants moved from aquatic to the terrestrial environments, new mechanisms were required for the exchange of nutrients - both photoassimilates from aerial organs to soil-anchored sections, and water and essential nutrients from the soil to the aerial, photosynthetic organs. Terrestrial plants developed complex vascular systems to provide, for example, roots with photoassimilates and to provide a photosynthetic organism with essential nutrients. Cells had to differentiate to acquire unique identities by establishing differential transcriptional networks. The vasculature serves both transport and communication between organs. Arabidopsis leaf veins are conjoint, collateral open and closed bundles, with xylem on the abaxial and phloem on the adaxial side^1^. The veins consist of at least seven different cell types with unique features identifiable by light and electron microscopy. In Arabidopsis, the abaxial phloem is composed of enucleate sieve elements (SE), as the actual conduits, which are coupled to companion cells (CC), and a third, poorly understood cell type, the phloem parenchyma (PP). When mature, the adaxial xylem consists of dead tracheary elements (TE), that are accompanied by xylem parenchyma (XP). The vascular parenchyma (VP) is often located at the interface between phloem and xylem. At earlier stages of development, xylem and phloem are separated by meristematic cells (procambium; open-type vasculature), that differentiate towards the tip of the leaf where procambium is absent (closed-type)^2^. Phloem and xylem exchange water and solutes in complex ways, and thus the cells must be equipped with specific sets of transporters. Moreover, it is likely that the different cell types have specialized metabolic activities. In addition, the vasculature plays important roles in communication by translocating hormones, small RNAs, and even proteins; and also appear to be involved in electrical signaling^3^.

PP is one of the poorly defined and least characterized vascular cell types. In Arabidopsis, PP has so-called cell wall ingrowths with a transfer cell appearance that are thought to play a role in amplifying the surface area to allow for higher transport rates^4,5^. Since in many plant species the interface between CC and SE (SE/CC) contains only few plasmodesmata, photoassimilate translocation requires an apoplasmic transport route^1^. SE/CC loading is mediated by the H^+^-sucrose symporter SUT1 (named SUC2 in Arabidopsis), which imports sucrose from the cell wall space^6^. We recently identified sucrose uniporters of the SWEET family in a specific subset of cells in the phloem, that likely represent PP^7^. SWEET expression in PP was confirmed using translational GFP fusions, using a new confocal microscopy method that enabled unambiguous PP identification^8^. A key goal of this work was to characterize the role of PP in more detail by identifying its mRNA outfit. This could serve, for example, as a basis for identifying transporters involved in phloem loading of other nutrients, as well as metabolic pathways active in these cells.

Understanding the complexity of plant cells at a single cell level has long been an active area of interest in plant biology. Comprehension at this level is crucial to determine subtle distinctions between, for example, the multitude of complex vascular cell types. Recently, high throughput single cell RNA-sequencing (scRNA-seq) has been used to provide new insights into root cell biology and development^9–13^. These latest technologies have allowed us to build on the resolution of prior atlases derived from the isolation of specific cell types, using fluorescent activated cell sorting of fluorescently-labelled protoplasts, and/or laser-capture microdissection^14,10,11^. scRNA-seq allows production of comprehensive data with higher resolution and with reduced complications and biases associated with the aforementioned techniques. To gain insights into the specific transcript profiles of the leaf vasculature, we optimized protoplast isolation protocols to enrich for leaf vascular cells. Using scRNA-seq, we identified unique and characteristic mRNA signatures, which revealed fundamental differences in amino acid metabolism and transport pathways in PP and CC important for phloem translocation as well as differences in hormone and glucosinolate metabolism.

## Results

### Enrichment of vascular protoplasts

Standard protoplast isolation protocols are efficient for protoplasting mesophyll cells for downstream applications such as transient expression. However, these procedures do not efficiently release cells from the vasculature. Here, we developed a methodology for isolating vascular-enriched protoplast populations from mature Arabidopsis leaves (Fig. 1, Supplementary Fig. 1). Initially, a fraction of the non-vascular cell types were removed using the ‘tape-sandwich’ method which effectively eliminated trichomes, guard cells, and epidermis from the abaxial leaf surface (Supplementary Fig. 1b)^15^. Cuts along the midvein further facilitated access of cell wall digesting enzymes to the vasculature (Fig. 1a, Supplementary Fig. 1b). Elevated concentrations of mannitol lead to an increased release of protoplasts (Fig. 1b, Supplementary Fig. 1c,d). We monitored the release by light microscopy, RT-qPCR, and fluorescence microscopy. The mRNA levels of marker genes were used to evaluate the enrichment of vascular protoplasts during optimization of the protocols (Fig. 1a-c). In addition, the release of intact vascular protoplasts was verified microscopically by monitoring fluorescently-labeled cells using stably transformed Arabidopsis lines expressing *pAtSWEET11:AtSWEET11-GFP*, a tentative marker for PP^7^, and *Q0990*, a procambial marker^16^ (Fig. 1d,e, Supplementary Fig. 1e. The bulk leaf (not subjected to protoplasting) transcriptome showed high correlation with bulk protoplast transcriptomes (Pearson’s correlation coefficient of 0.9) (Supplementary Fig. 2a, Supplementary Table 1). The high correlation indicates that the protoplasting protocol used here did not impact the relative abundance of cells in the leaf and supported the notion that the scRNA-seq data derived from the protoplasts should cover essentially all cell types present in a mature leaf (the term enriched here thus refers to enrichment over standard protocols).

**Fig. 1:**
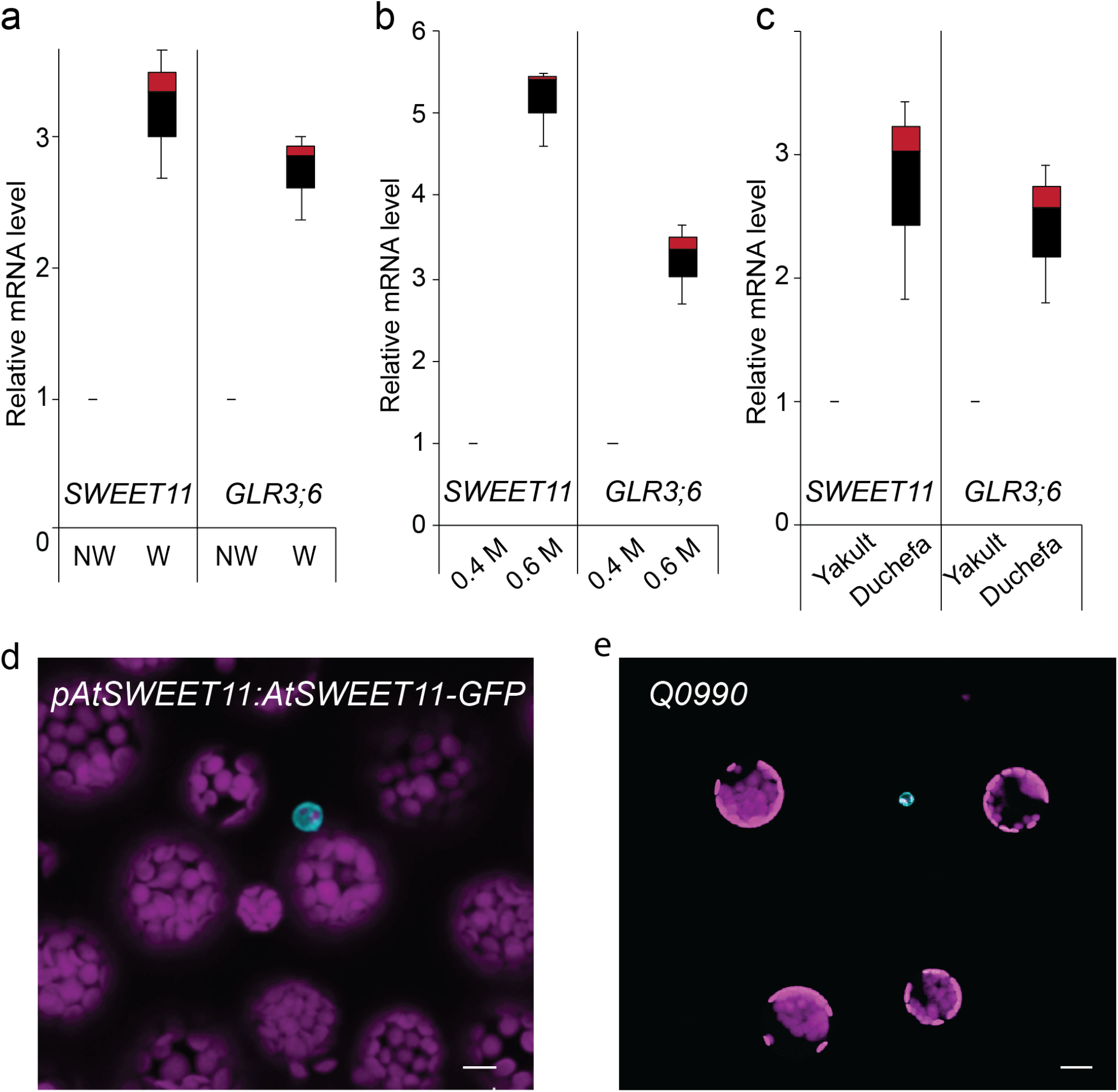
Strategies to enrich vascular protoplasts. **a-c**, Box plot representation of the RT-qPCR analysis results showing the relative transcript level of the marker for the phloem parenchyma, *SWEET11*, and the marker for xylem parenchyma, *GLR3;6. UBQ10* was used as an internal control for normalization. The abaxial epidermis was removed from leaves harvested from 6-weeks-old plants grown under short day conditions and was treated as indicated. The data shown are from a single experiment with three technical replicates (mean ± SE, n = 3). **a**, Leaves from which the abaxial epidermis had been stripped were either cut on each side of the main vein before enzymatic digestion of the cell wall (wounded, W) or directly placed in the digesting enzyme solution (nonwounded, NW). **b**, Leaves from which the abaxial epidermis had been stripped were cut on each side of the main vein and incubated in the enzyme solution containing 0.4 M or 0.6 M mannitol. **c**, Protoplasts isolated from leaf sample prepared as in **b** were incubated in an enzyme solution composed of cell wall degrading enzymes (cellulase Onozuka R-10, macerozyme R-10) obtained from Yakult (Tokyo) and Duchefa (Haarlem). **d**, GFP fluorescence (cyan) marking phloem parenchyma cells in leaf protoplasts. Protoplasts were isolated from 6-week-old *pAtSWEET11:AtSWEET11-GFP* plants expressing a phloem parenchyma cell-specific marker. Magenta, chlorophyll autofluorescence. Scale bar: 10 μm. e, GFP fluorescence (cyan) marking procambium cells in leaf protoplasts. Protoplasts were isolated from 6-week-old *Q0990* plants expressing a procambium cell-specific marker. Magenta, chlorophyll autofluorescence. Scale bar: 20 μm.

### Single cell RNA-sequencing of vascular-enriched leaf protoplast population

scRNA-seq libraries were produced from vasculature-enriched leaf protoplast populations, with 5,230 single-cell transcriptomes obtained from two biological replicates. Sequencing to a depth of ∼96,000 reads per cell was undertaken, and identified a median number of 3,342 genes, and 27,159 unique molecular identifiers (UMIs, representing unique transcripts), per cell (Supplementary Fig. 2b). Unsupervised clustering using Seurat^17^ identified 19 distinct cell clusters (Fig. 2a, Supplementary Video 1, Supplementary Table 2). Plotting the transcriptomes from the two replicates in two dimensions using Uniform Manifold Approximation and Projection (UMAP)^18^ revealed an overlapping distributions of cells, and a similar proportion of cell identities (Supplementary Fig. 2c). mRNA profiles and relative cell numbers in the clusters were highly correlated in the replicates (Supplementary Table 2). To assign cell identity to the clusters, we examined the specificity of transcripts of known marker genes (Fig. 3, Supplementary Fig. 3, Supplementary Table 3 and 4).

**Fig. 2:**
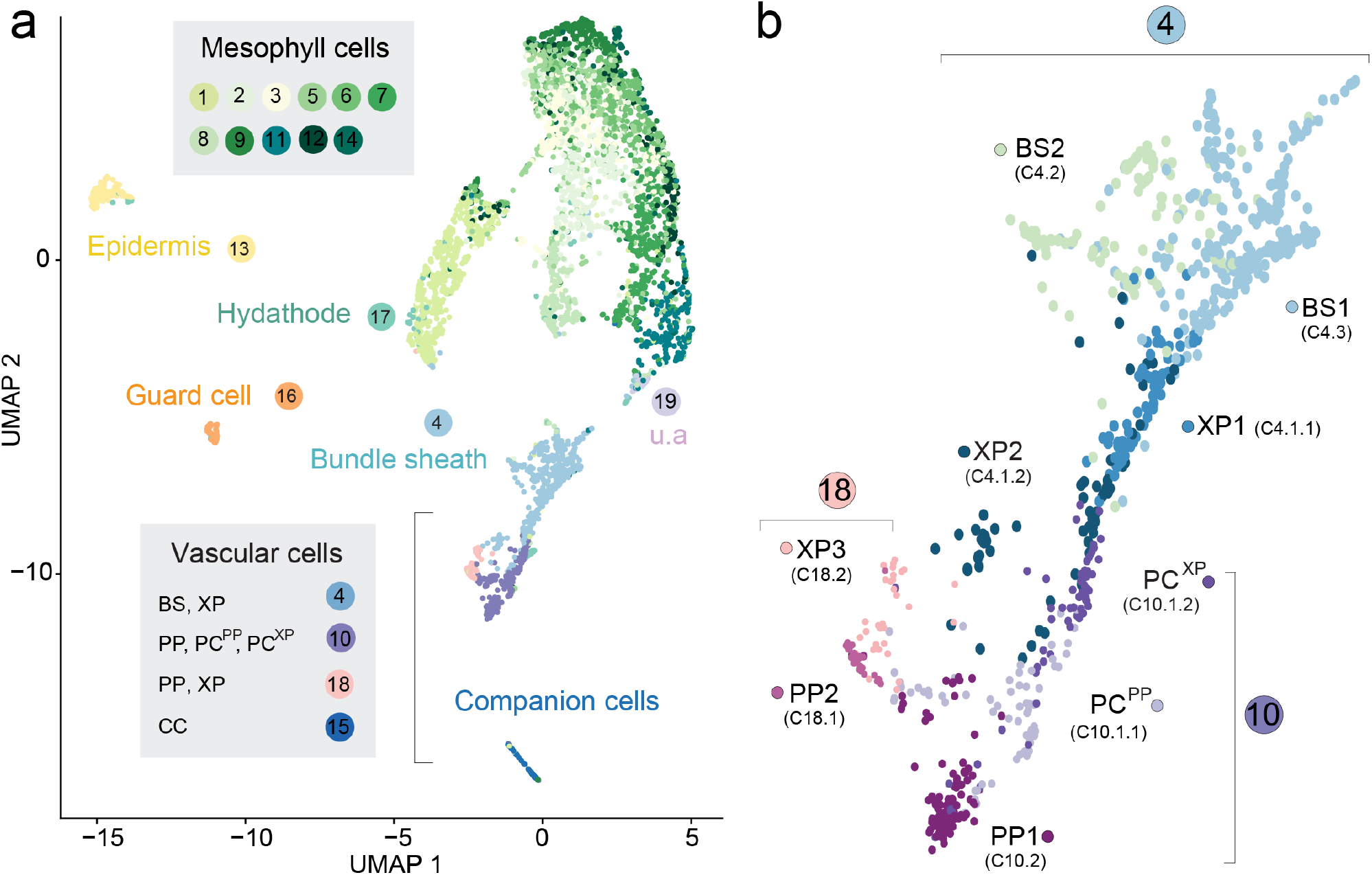
Assignment of cellular identity to clusters. **a**, UMAP dimensional reduction projection of 5,230 Arabidopsis leaf cells. Cells were grouped into 19 distinct clusters using Seurat^65^. The cluster number is shown and colored based on the colors assigned to each cell type (u.a. – unassigned, i.e., cluster could not be assigned to a known cell type). Each dot indicates individual cells colored according to the cell type assigned. **b**, Magnification of subclusters C4, C18, and C10. Different colors indicate distinct cell identities. PC: procambium, BS: bundle sheath, XP: xylem parenchyma, PP: phloem parenchyma, PC^XP^: procambium cells with features relating to xylem differentiation, PC^PP^: procambium cells with features relating to phloem differentiation.

**Fig. 3:**
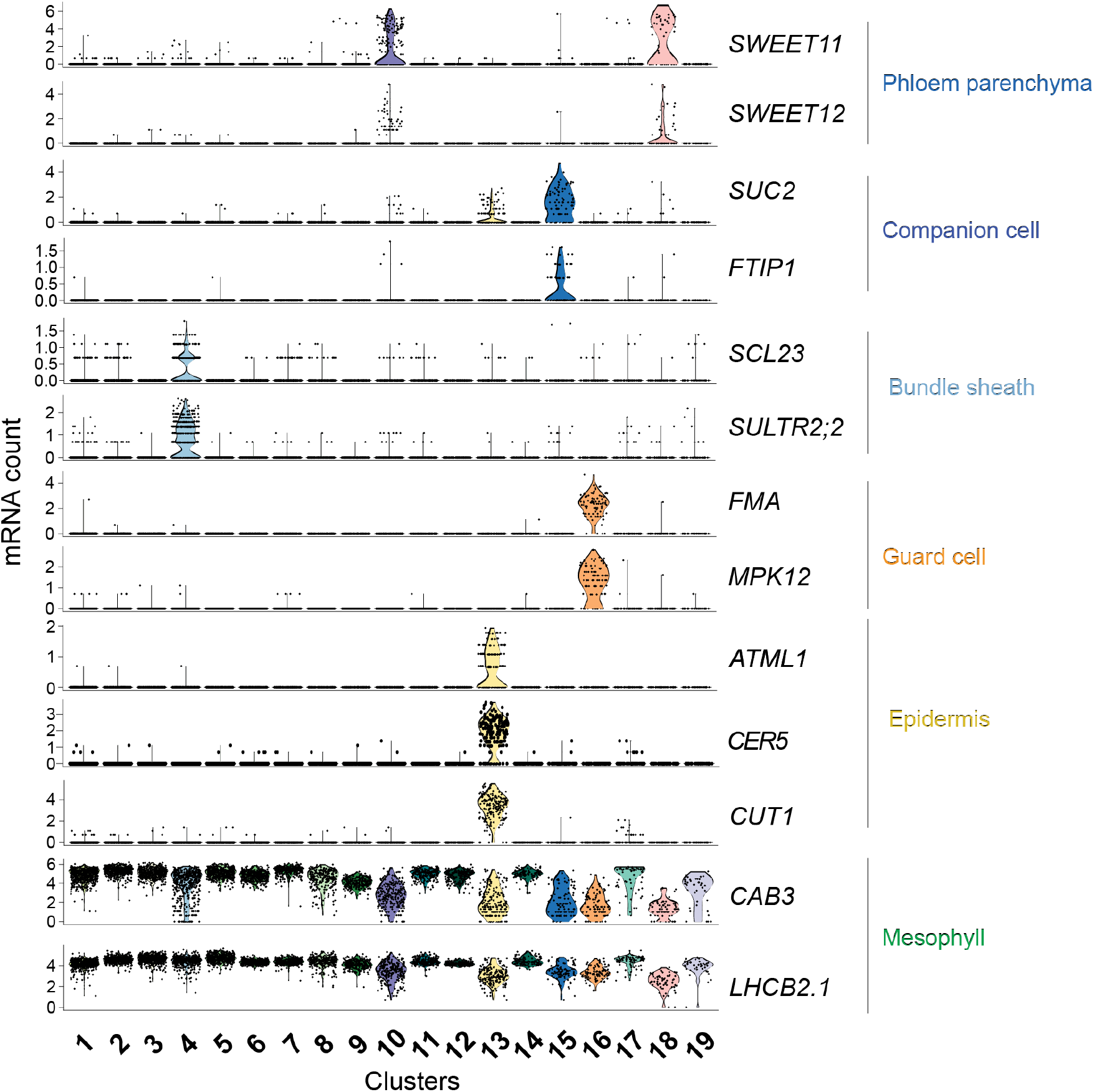
mRNA levels of marker genes in clusters used to assign cell types. Violin plots showing transcript enrichment of known cell type-specific marker genes across clusters. Clusters are indicated on the x-axis. The name of the cell type assigned to each cluster is indicated on the right side of the violin plots.

The 19 clusters cover all major cell types of the leaf. The population was dominated by mesophyll cells (∼75% of all cells). as indicated by the enrichment of known mesophyll markers (*CAB3, LHCB2*.*1*, and *CA1*)^19,20^ (Fig. 3, Supplementary Fig. 3a). The mesophyll group comprised twelve clusters (Clusters, or C1, 2, 3, 5, 6-9, 11, 12, 14; Fig. 2a, Fig. 3; Supplementary Table 4) and formed three major branches on the UMAP plot. Naively, two separate mesophyll clusters can be expected, one for palisade and one for spongy parenchyma. *FIL* (*YAB1*) transcripts, markers for spongy parenchyma were enriched in C2 and C7, possibly indicating that these clusters may represent spongy parenchyma (Supplementary Fig. 3a)^21^. Among the mesophyll clusters, C6 and C9 were enriched for rRNAs (Supplementary Table 4). We did not remove rRNA before sample preparation and did not filter cells that showed higher rRNA transcript levels since rRNA levels can vary between cells. Whether C6 and 9 represent unique biologically relevant cell populations in the leaf or are due to artifacts was not further evaluated. While the mesophyll clusters likely contain interesting information, we did not characterize them further, as we decided to focus on the vasculature.

Beyond the mesophyll, epidermal cells were identified in C13, based on enrichment of the epidermis-specific transcription factor *AtML1*^22^ and wax-related genes (Fig. 3). Epidermal guard cells were found in C16, indicated by the specific enrichment of *FAMA* and guard cell-specific MAP kinases (Fig. 3, Supplementary Fig. 3a). Enrichment of transcripts of the purine transporter *PUP1* and the homolog of carrot chitinase, *EP3*, indicated that C17 comprises hydathodes cells (Supplementary Fig. 3a)^23^.

Several clusters showed a vasculature identity, based on the enrichment of known marker genes, indicating that the enrichment for these cell types was successful. The vasculature is composed of phloem and xylem, often separated by a meristematic layer, the procambium, which initiates and maintains vascular organization. We assigned clusters to the seven known cell types present in leaves (Fig. 2). Additionally, we identified cell types with identities that had properties of two cell types, e.g. xylem and procambium.

Cluster 4 was found to contain bundle sheath (BS) and xylem cells and was subsequently subclustered into three cell populations, C4.1, C4.2, and C4.3 (Fig. 2b, Supplementary Table 5). Subclusters C4.2 and C4.3 were enriched for the BS marker *SCL23*, and the sulfate transporter, *SULTR2;2* (Fig. 2b, Fig. 3, Supplementary Fig. 3b,c). C4.2 was enriched for genes involved in photosynthetic processes, possibly indicating a differentiation of BS cells (Supplementary Table 5). The existence of two BS clusters is consistent with morphological descriptions^24^. C4.1, C10, C15, and C18, with a total of 478 cells, were assigned as vascular cell types (Fig. 2, Fig. 3, Supplementary Fig. 3a, Supplementary Table 3).

### Degree of kinship relations among BS, XP, PC and PP

Clusters composed of BS, xylem parenchyma (XP) and procambial (PC), and PP cells show a J-shaped relatedness in our UMAP plot (Fig. 2). BS is related to two types of xylem cells (XP1 and XP2), that abut the PC. PC subclusters into two subsets closely related to XP1 (PC^XP^) and PP1 (PC^PP^), respectively. The PC^PP^ is adjacent to the two PP clusters PP1 and PP2. XP3 and PP2 form a separate group, which together with XP2, form the hook of the J shape in our UMAP plot (Fig. 2b).

### Xylem-related clusters

The enrichment of *GLR3;6*^3^ and *ACL5* markers identified C4.1 as parenchymatic xylem cells (Supplementary Fig. 3a,d-i). Further clustering of C4.1 resulted in two subgroups, XP1 and XP2, distinguished by the relative enrichment of photosynthetic genes (Supplementary Table 5). An additional subcluster with xylem identity, XP3, was observed as a subgroup of cells from C18. XP3 was enriched with xylem markers as well as those for vascular parenchyma^25^ (Supplementary Fig. 3d-i, Supplementary Fig. 4). Transport proteins were enriched in the xylem clusters (XP1-XP3), as one may expect from their role in supplying tracheal elements with ions and nutrients for xylem translocation. These included amino acid transporters, such as the H^+^/amino acid symporter *AAP6* and the UmamiT amino acid exporter family member *UmamiT22* (refs.^26–28^). The amino acid export protein *GDU4* was also enriched in the xylem clusters. Multiple SULTR low-affinity sulfate transporters, which play pivotal roles in sulfate transport in xylem parenchyma cells, were detected in the xylem cells as well as the boron transporter *NIP6*.*1*, the glucosinolate importer *GTR2/NPF2*.*11*^29,30^, the heavy metal transporter *HMA2*, and the plastidic bile acid transporter *BAT5*/*BASS5* (Supplementary Table 6).

### Phloem-and xylem-related subpopulations of the procambial subclusters

Arabidopsis procambium constitutes a bifacial stem cell population producing xylem on the adaxial pole and phloem on the abaxial pole^32,33^. On our UMAP plot, the XP clusters are in proximity to Cluster 10. Cluster 10 could be further subdivided into three subclusters: the PP1 (C10.2) and two distinct procambial populations, C10.1.1 and C10.1.2 (Fig. 2, Fig. 4a, Supplementary Tables 7 and 8). While cells in PC^PP^ corresponds to procambial cells involved in the maintenance of meristematic identity and the differentiation of phloem cells, cells of PC^XP^ appear to be more closely related to the formation of xylem cells, and is located closer to the XP clusters (Fig. 4, Supplementary Tables 7 and 8).

**Fig. 4:**
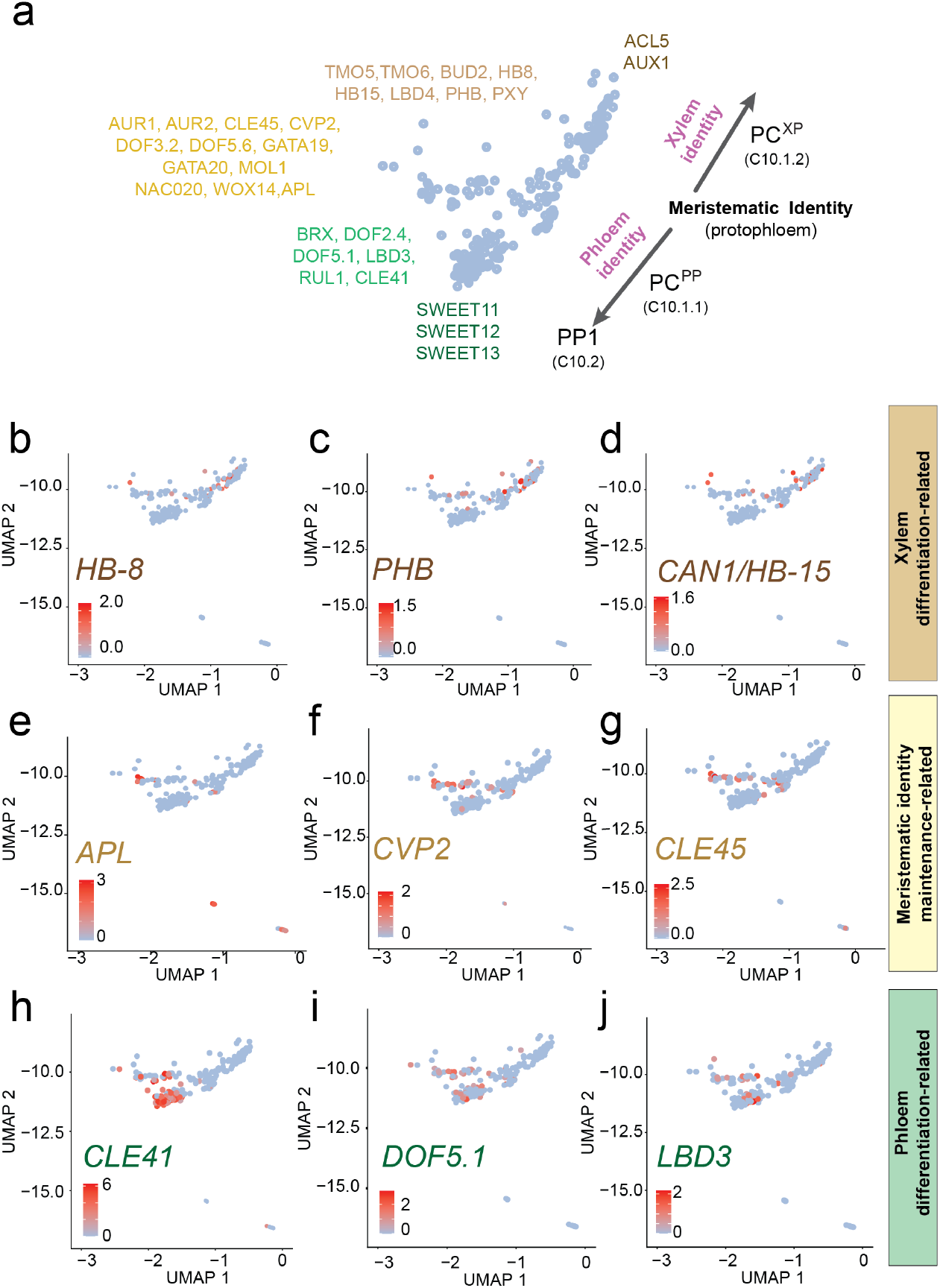
Identification of the procambium cell cluster with distinct procambium cell identities. **a**, Schematics representing subpopulations of Cluster 10. Genes enriched in the subpopulations are indicated. **b-c**, UMAP showing enrichment of transcripts of genes related to xylem differentiation. **e-g**, UMAP showing enrichment of transcripts of genes related to maintenance of protophloem pluripotency and differentiation. **h-j**, UMAP showing the distribution transcripts related to phloem differentiation.

Cells in the PC^XP^ cluster were enriched with transcripts for homeodomain leucine-zipper (HD-Zip) transcription factor, *HOMEOBOX GENE 8* (*HB-8*)^33^ transcripts, known to trigger xylem differentiation (Fig. 4b). Adaxial HD-Zip transcription factors, such as *REVOLUTA/INTERFASCICULAR FIBERLESS1* (*REV/IFL1*), *PHABULOSA* (*PHB*), and *CORONA* (*CNA/AtHB15*), were also enriched in this cluster (Fig. 4c,d, Supplementary Fig. 5). Conversely, the PC^PP^ cluster closest to the PC^XP^ was enriched with known factors involved in the maintenance of protophloem identity and pluripotency, such as *COTYLEDON VASCULAR PATTERN 2* (*CVP2*)^34^ and *CLAVATA3/EMBRYO SURROUNDING REGION-RELATED 45 (CLE45)*^35,36^ (Fig. 4e-g, Supplementary Fig. 5). *ALTERED PHLOEM DEVELOPMENT* (*APL*), a major phloem marker which inhibits xylem differentiation, was also detected in this region^37^ (Fig. 4e). Transcripts related to phloem differentiation, including *CLE41, DOF5*.*1* and *LBD3* were enriched in the lower region of PC^PP^ (Fig. 4h-j, Supplementary Fig. 5). Strikingly, the arrangement in the UMAP plot, of the PC^PP^ cluster facing the PP cluster (C10.2) and the PC^XP^ facing the XP clusters, is analogous to the morphological positioning. A detailed description of the procambium clusters is included in the Supplementary Text.

### Two phloem parenchyma clusters

SWEET11 and SWEET12 efflux transporters had previously been reported to be expressed in the PP^8,38^. Sucrose released by the SWEETs into the apoplasm is taken up into the neighboring SE/CC via the sucrose/H^+^ symporter SUC2/SUT1^6,39^. Interestingly, we identified two clusters, C10.2 (PP1) and C18.1 (PP2), specifically enriched for *SWEET11* and *SWEET12* (Fig. 3, Fig. 5a-f). Cells in the two PP clusters are enriched for transcripts related to transport processes, reflecting the roles of PP as a critical cell type required for loading diverse substrates into the phloem (Supplementary Tables 7, 9-11). We identified an additional member of the Clade III SWEET family, *SWEET13*, in the cells that expressed *SWEET11* and *SWEET12* (Fig. 5g-i). *SWEET13* transcripts show compensatory accumulation in *sweet11;sweet12* double knockout mutants, indicating redundant roles with SWEET11 and SWEET12^38^. Using a modified protocol from a recently developed confocal imaging method that enables identification of PP ^8^, we were able to validate the presence of SWEET11 and SWEET13 in the PP-cells in *pSWEET13:SWEET13-YFP* (Fig. 5p) and *pSWEET11:SWEET11-2A-GFP* lines (Fig. 5q,r).

**Fig. 5:**
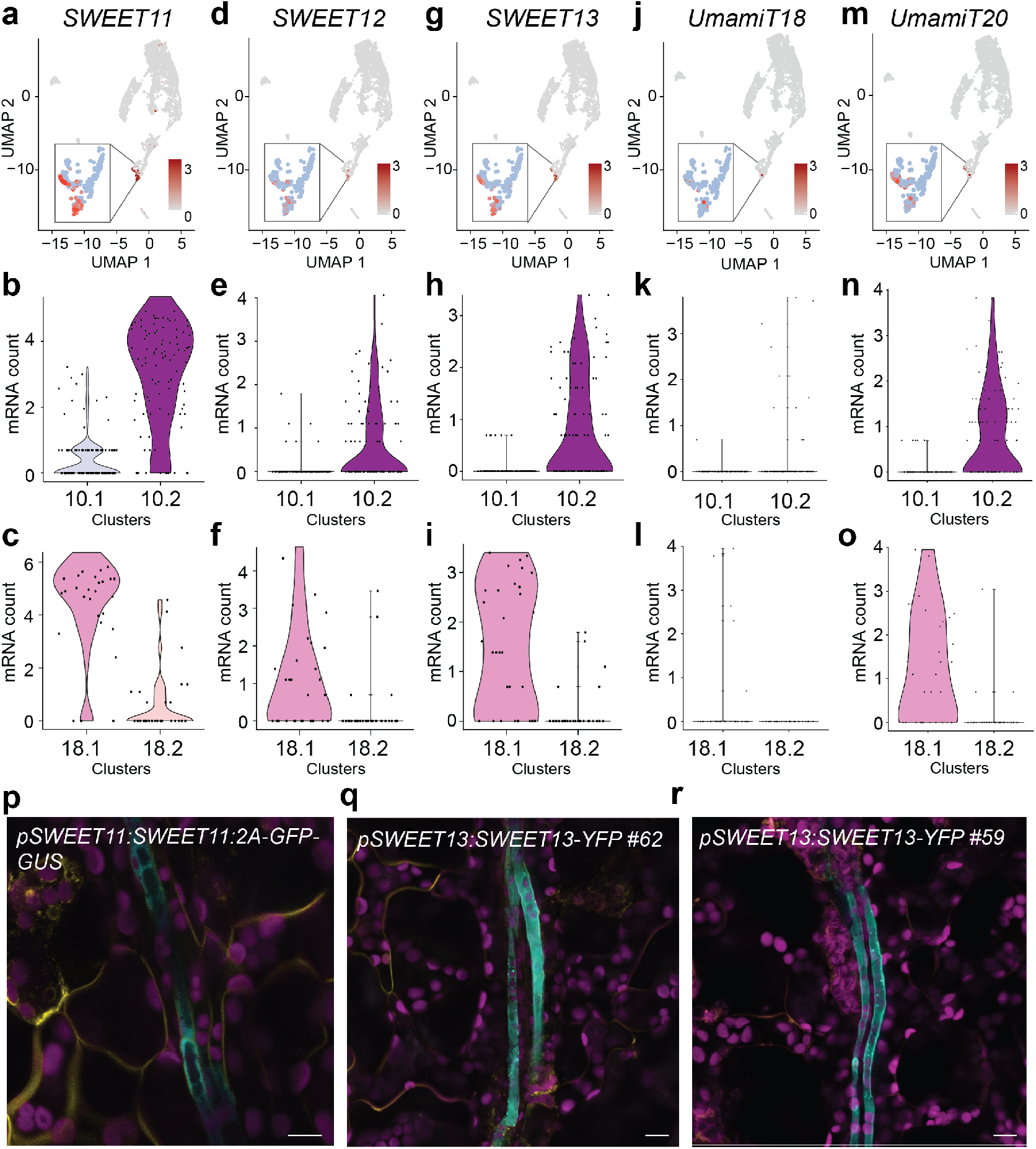
Three SWEET sucrose transporters and UmamiT amino acid transporters mark the phloem parenchyma cluster. **a-o**, UMAP and violin plots of C10 and C18 subclusters showing enrichment of *SWEET11*(**a-c**), *SWEET12*(**d-f**), *SWEET13*(**g-i**), *UmamiT18/SIAR1*(**j-l**), and *UmamiT20*(**m-o**) transcripts in phloem parenchyma clusters. Subcluster 10.1 corresponds to PC, 18.2 to XP3, and 10.2 and 18.1 to PP. Inset show magnification of C10 and C18. **p-r**, Confocal microscopy images of *SWEET11:SWEET11-2A-GFP-GUS* and (**p**), *pSWEET13:SWEET13-YFP*(**q**,**r**) transgenic plants showing specific GFP(**p**) or YFP(**q**,**r**) signal in the PP. Magenta, chlorophyll autofluorescence. Yellow, FM4-64FX. Cyan GFP fluorescence (**p**) or YFP fluorescence(**q**,**r**). Scale bars: 10 μm.

UmamiT family members have been described as cellular exporters for amino acids^40^. We thus speculated that UmamiTs may play analogous roles for the efflux of amino acids from PP as the SWEETs do for sucrose. Consistent with our hypothesis, we found *UmamiT18/SIAR1* mRNA in PP (Figs 5j-l). Vascular expression in leaves is supported by *UmamiT18/SIAR1* transcriptional fusion lines^40^. In addition, transcripts for six other members of the UmamiT family (UmamiT12, 17, 20, 21, 28, 30) were enriched in the two PP clusters (Fig, 5m-o, Supplementary Fig. 6a-f, Supplementary Tables 7 and 11). Notably many of the PP-specific *UmamiT*s were coexpressed with each other as well as *SWEET11* and *12* (Supplementary Fig. 6g, Supplementary Table 12).

As many transcripts related to transport processes were specifically enriched in the PP clusters, and many of them were coregulated (Supplementary Table 12), this may indicate that they are subject to control by the same transcriptional networks. We therefore searched for transcription factors that might be responsible for coregulation of the functionally related transporters. We identified multiple leucine zipper genes (bZIP) either specific to the PP clusters (*bZIP*6, *bZIP7, bZIP9*), or present in PP and other clusters (*TGA7, bZIP11*; Supplementary Fig. 7). We were able to confirm that *bZIP9* promoter activity was specific for the PP cells in the leaf vasculature, using a transcriptional fusion, *pbZIP9:GFP-GUS* (Fig. 6a-d). Strong GUS activity was also detected in cell types known to be involved in apoplasmic transport steps, such as the unloading zones of anthers and seeds, and the veins of petals, receptacles, ovules, transmitting tract, and funiculi (Supplementary Fig. 8). The cell specificity of bZIP9 is consistent with a possible role in the activation of target genes involved in transport process across different tissues. The nature of the separation of PP into two subclusters will require further analysis.

**Fig. 6:**
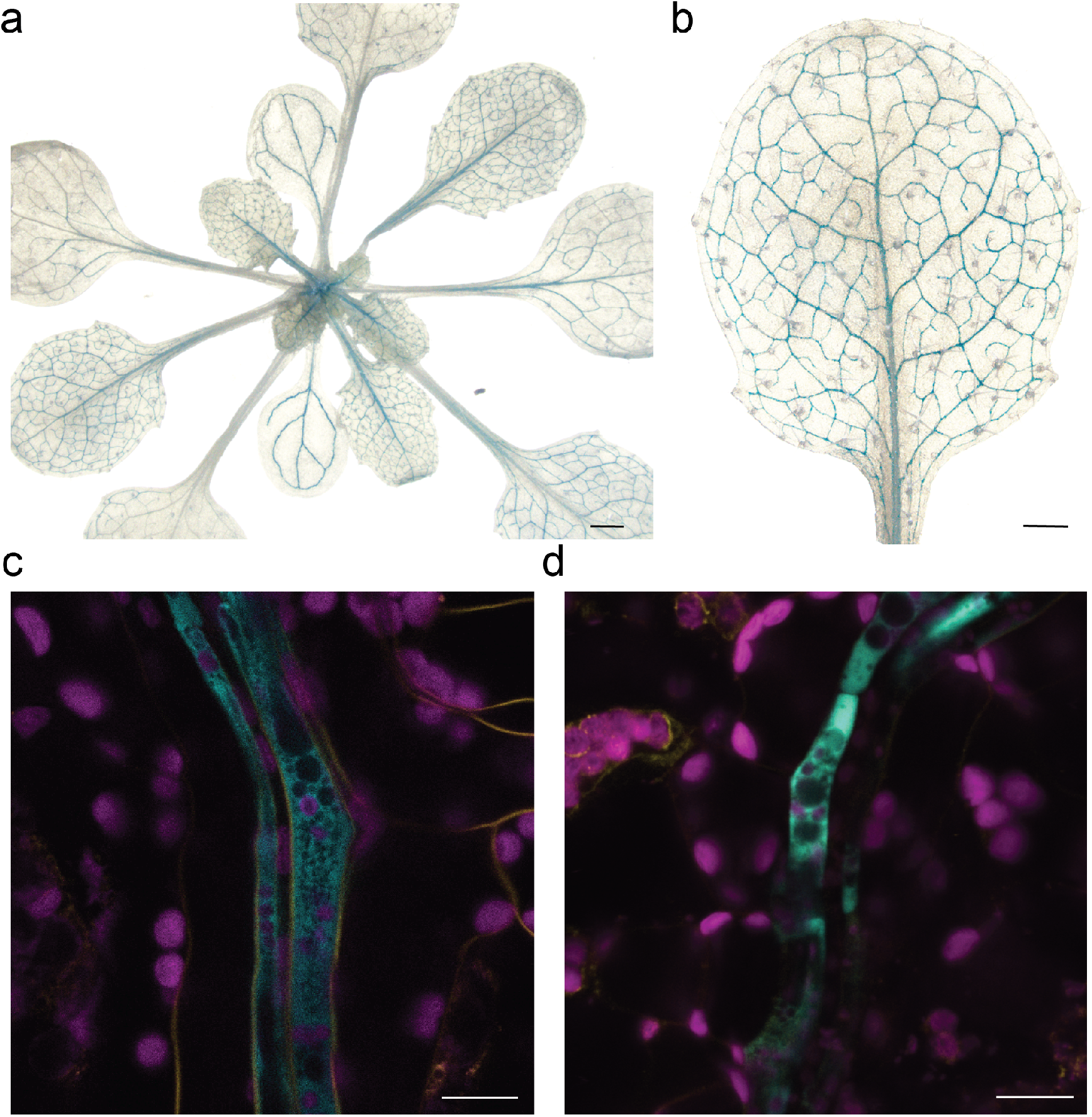
Reporter gene analysis of *pbZIP9:GFP-GUS* plants. **a**,**b**, GUS stained transgenic plants expressing the transcriptional *pbZIP9:GFP-GUS* reporter construct show GUS activity in the leaf vasculature. Scale bars: 1 mm (**a)** and 0.5 mm (**b). c**,**d** Confocal microscopy images of *pbZIP9:GFP-GUS* reporter lines showing GFP signal specific in the phloem parenchyma cells. Magenta, chlorophyll autofluorescence. Yellow, FM4-64FX, Cyan, GFP fluorescence. Scale bar: 10 μm.

### The companion cell cluster

Companion cells, which acquire carbon and nitrogen from the adjacent PP, but also maintain the functionality of the enucleate sieve elements, cluster separately from all other cell types. C15 was designated as CC based on the enrichment of transcripts for the sucrose/H^+^ symporter *SUC2/SUT1*, the H^+^-ATPase *AHA3*, and genes such as *FTIP* and *APL* (Fig. 3, Supplementary Fig. 3a). CCs were enriched for transcripts of various transporters, including the amino acid/H^+^ symporter *AAP2* (ref.^41^), the potassium channel *KAT1* (ref.^42^), the hexose uniporters *SWEET1* and *SWEET4* (ref.^43^), and the tonoplast peptide transporter *PTR4/NPF8*.*4* (ref.^44^) (Supplementary Table 11). All these markers are also enriched in the CC translatome^45^. Notably, many genes involved in control of flowering were also enriched in CCs (Supplementary Table 3).

### PP and CC cells undergo differentiation through distinct pathways

The clear separation of PP and CC in the UMAP was striking and indicates the presence of distinct differentiation <u>pathways</u>. The MYB coiled-coil-type transcription factor APL plays a key role for the definition of phloem identity^37^. In our datset, APL was preferentially expressed in CC, but also detected in the PC^PP^ cluster (Fig 4e and Supplementary Fig. 3a). As APL function is necessary for the development of the SECC complex, one may propose two hypotheses: (i) APL serves as a master regulator of all phloem cells, or (ii) PP develops independently of the SECC differentiation. While *apl* mutants are characterized by drastic reduction of CC marker *SUC2* ^37^, *apl* retains expression of multiple PP markers (Fig. 7b); PP marker transcripts were even > fourfold higher in the *apl* mutant (Fig. 7a). Whether the increase of PP-marker transcripts in *apl* is due to a compensatory mechanism in which APL represses PP differentiation, as suggested for xylem differentiation^37^, remains to be elucidated.

**Figure 7.**
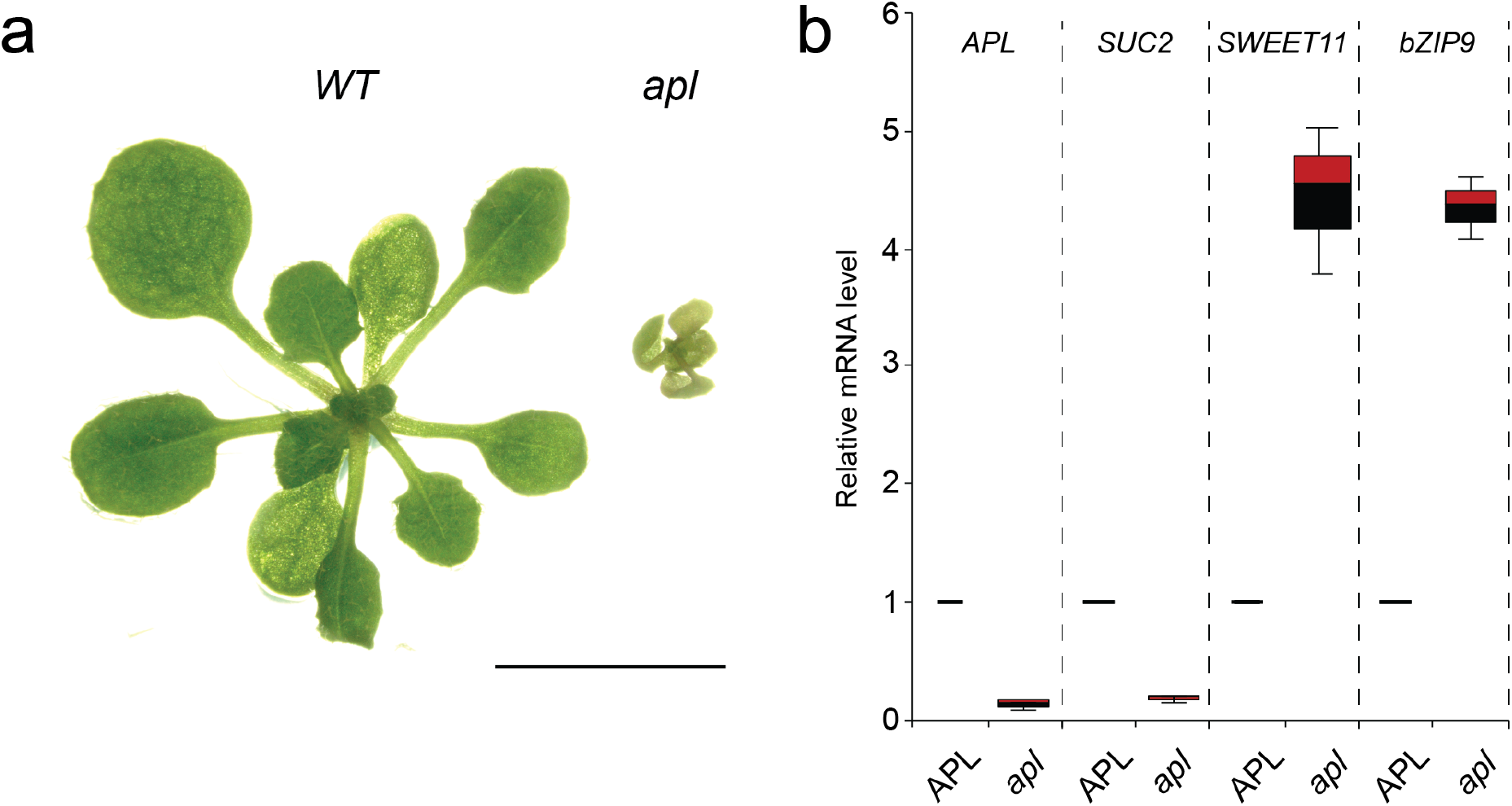
Distinct PP and CC marker gene expression in the *apl* mutant. a, Morphology of *apl* mutant (right) grown on LD conditions for 2 weeks. WT plant grown under the same condition is shown on the left side. Scale bar: 1 cm b. RT-qPCR analysis of CC-marker gene (*SUC2*), and PP-marker genes (*SWEET11* and *bZIP9*). Segregating seeds from heterozygous parents were plated on MS for 2 weeks. The first and second leaves from plants homozygous for the APL mutation (*apl*) and heterozygous for the mutation or WT (APL) were collected for RNA extraction and RT-qPCR. Three independent replicates showed similar results and a representative experiment with three technical replicates are shown (mean ± SE, n = 3).

### Unique and complementary metabolic landscapes of PP and CC

Amino acid transporters of the AAP and UmamiT families are relatively non-selective and transport many of the proteogenic and other amino acids^28,46^. One may therefore assume that the translocation of most amino acids is similar and that relative amino acid levels are mainly determined by the relative rates of biosynthesis, that the amino acids all enter the translocation stream in similar ways and that the relative levels do not change substantially during translocation. However, labeling studies have shown that amino acids behave very differently, with some being effectively metabolized along the path of source-to-sink partitioning, while other stay largely unmetabolized^47^. Surprisingly, the amino acid distribution in stems is also varying for different amino acids^48^. These phenomena might be explained by differential distribution of metabolic activities in different vascular cell types. We therefore carried out a pathway activity analysis to define a pathway activity score (PAS) that quantifies the average expression of the pathway’s genes in one cell type relative to the other cell types^49^. The PAS values of the CC and PP were strikingly different for both amino acid biosynthetic and degradation pathways (Fig. 8). While the activity of biosynthetic amino acid pathways was low in CC, PP-containing clusters showed elevated metabolic activity. A comprehensive analysis of pathway activities across all clusters revealed many other interesting features unique to specific cell types. For instance, the epidermis cluster (C13) was enriched with biosynthetic activities for wax esters, cuticular wax, the suberin monomer and very long chain fatty acids. These substances function as coatings in the epidermis, serving as a moisture barrier and protecting plants from pathogens. The PP-including cluster, C18, showed high activity of hormone pathways such as ABA, ethylene, JA, and GA, and the BS and XP clusters (C4) and the two PP-including clusters (C10 and C18) were enriched for glucosinolate biosynthesis activity (Supplementary Text, Supplementary Figs. 9 and 10). C19 may represent so-called S-cells, cell likely involved in defense responses, based on the enrichment of programmed cell death pathways and PAS values for insect chewing-induced glucosinolate breakdown^50^ (Supplementary Table 13, Supplementary Text). The two PP clusters (C10, C18) showed high PAS values for callose biosynthesis, consistent with callose deposition in PP transfer cells^51^. We also provide a PAS analysis across all cell types that is not discussed further here (Supplementary Fig 10, Supplementary Text).

**Fig. 8.**
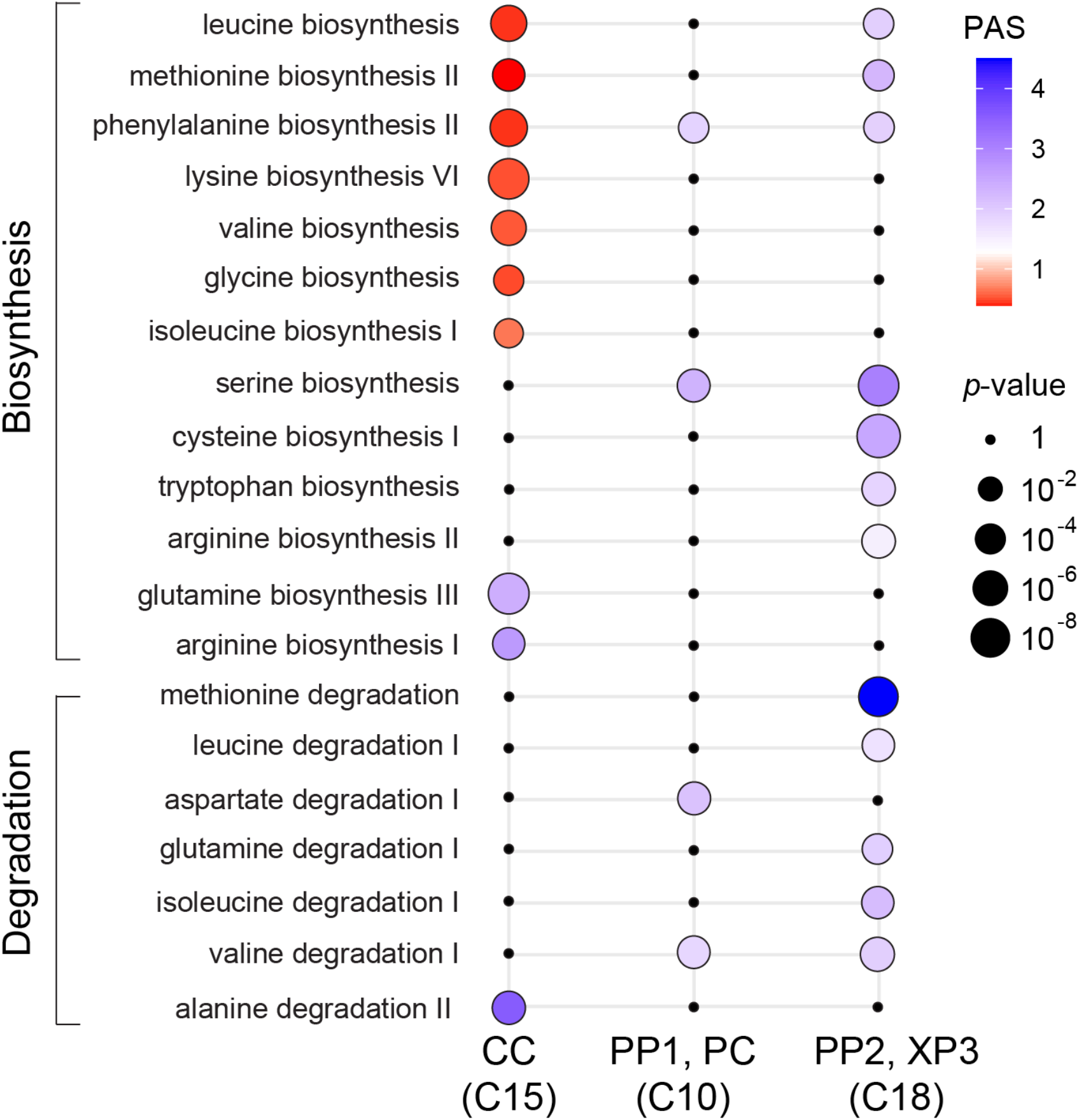
Amino acid biosynthesis and degradation pathways are differentially represented in the CC PP, PC, and XP3 cells. Metabolic pathway activities of amino acid biosynthesis and degradation pathways in Clusters 15, 10, and 18. Statistical significance is represented as differences in dot size. Statistically insignificant values are shown as black dots (random permutation test, *p* >0.05). Colors represent the pathway activity score (PAS); a score <1 (red) reflects a lower than average activity of the pathway in the given cell type, a score >1 (violet) indicates a higher activity. Activities were compared between all clusters in the scRNA-seq dataset.

### Cell-type specific expression of plasmodesmatal proteins

Plasmodesmata (PD) are highly complex channels that interconnect plant cells, likely transporting solutes, metabolites, small RNAs and proteins. Different cell types have unique types of plasmodesmata, including that which connects CC to enucleate sieve elements to supply all components necessary for function. The PDs typically are branched on the CC side and have a single pore on the sieve element side. We here found that transcripts of PD genes such as the *PLAMODESMATA-LOCATED PROTEIN* (*PDLP*)s and *MULTIPLE C2 DOMAINS AND TRNAMEMBRANE REGION PROTEIN (MCTP*)s showed distinct expression patterns in different cell types (Fig. 9a-d, Supplementary Fig. 11a,b)^52,53^. For example, *PDLP6, 7*, and *8* transcripts were found in PP and CC, while *PDLP2* and *3* transcripts were present in mesophyll cells (Fig. 9a-d, Supplementary Fig. 11a). The cell-type specificity of *PDLP*s was correlated with their phylogenetic relationships (Fig. 9e). Our results are consistent with data from GUS reporter fusions for the *PDLP7* and *8* promoters^54^. *MCTP1* and *3* transcripts were mainly enriched in the vasculature (Supplementary Fig. 11b). In particular, *MCTP1* transcript was specifically enriched in CC, consistent with a role in trafficking the florigen protein FLOWERING LOCUS T (FT), from companion cell to sieve elements^55^. In comparison, *MCTP4, MCTP6* and *MCTP15* are ubiquitously expressed in vascular and mesophyll cells (Supplementary Fig. 11b). Overall, our data supports the concept of PD-type specific assembly of PDs.

**Fig. 9:**
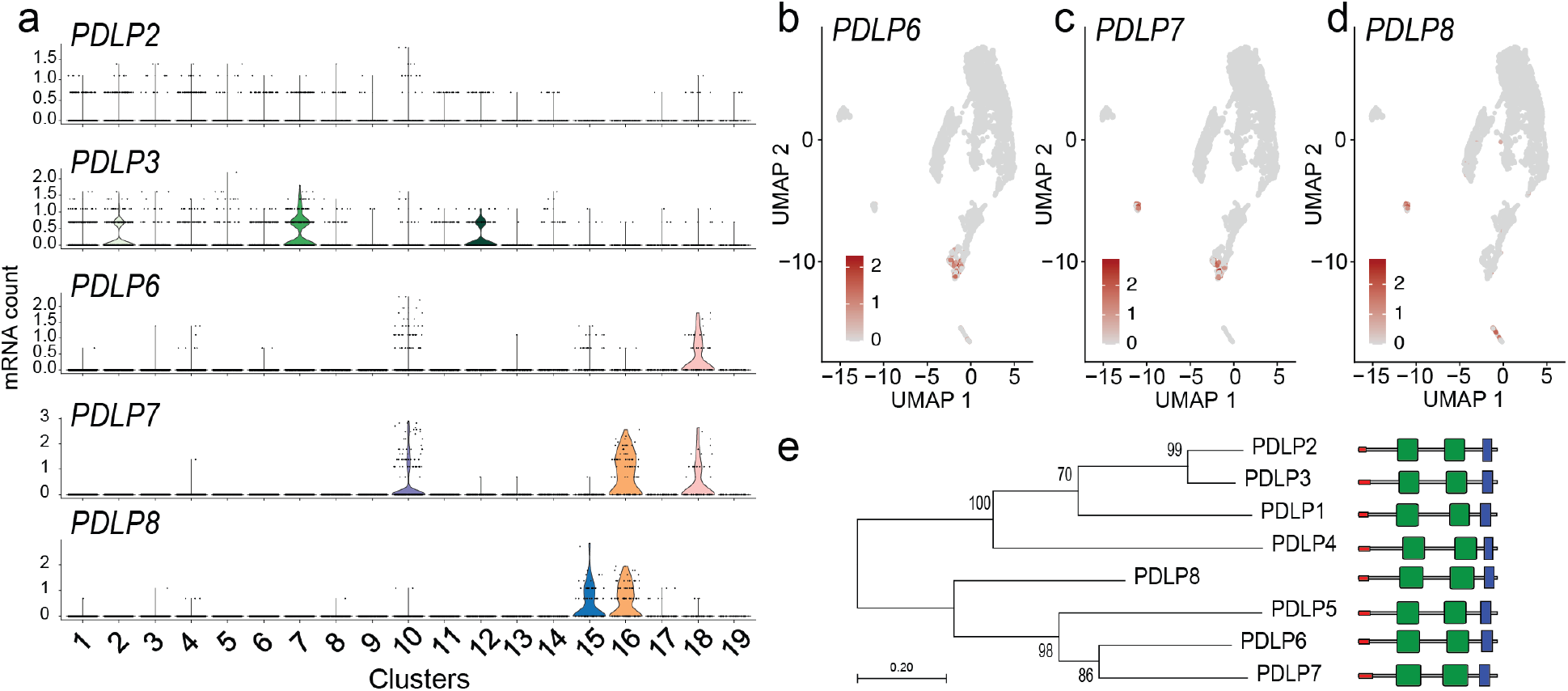
Transcript enrichment of *PDLP* genes in vascular cell types. **a**, Violin plot showing transcript enrichment of *PDLP*s. **b-d**, UMAP plot showing the enrichment of *PDLP6* **(b)**, *PDLP7* **(c)** and *PDLP8* **(d)** transcripts in the PP clusters (**b**,**c**), guard cell cluster (**c**,**d**), and CC cluster (**d**). **e**, Phylogenetic analysis of PDLPs in Arabidopsis. The phylogenetic tree was generated with the maximum likelihood method implemented in PhyML^72^. Percent support values from 1000 bootstrap samples are shown. Protein motifs predictions are based on the SMART database (http://smart.embl-heidelberg.de). DUF26 domains, transmembrane region and signal peptide are shown in green, blue and red, respectively.

## Discussion

Despite the advances in scRNA-seq technologies, application to plant cells still faces the challenge of removing cell walls to allow the release of individual cells and the penetration of the buffers into cells. Recent studies (using Arabidopsis, rice, as well as maize aerial tissues) clearly demonstrate the difficulty of capturing vascular cell types. For example, a key cell type in the vasculature, the PP, was not identified in any of the recently published scRNA-seq data from roots^9–13^. Here, we systematically optimized protoplast isolation protocols to enrich vascular cell types and produced a single cell transcriptome and metabolic activity score atlas that covers essentially all known cell types in the Arabidopsis leaf. The scRNA-seq confirmed that except for trichomes, abaxial epidermis and guard cells which were intentionally removed, all known nucleus-containing cells of the leaf were represented in the dataset. In particular, all cells from the leaf vasculature were identified.

The vascular cells had identities clearly distinct from those of the epidermis, guard cells and mesophyll. The bundle sheath formed a supercluster, together with all vascular cells including the vascular meristem, except the companion cells (CC). A major finding within this atlas was that the kinship relation of vascular cells was highly similar to the actual morphology of the vasculature, with the exception of the CC, which formed a unique island separate from BS, xylem, procambium and the phloem parenchyma. The arrangement of the XP-PC^XP^-PC^PP^-PP clusters in the UMAP plot reflected a potential developmental trajectory, in which the meristematic cells are localized in the center of the cluster and the surrounding cells differentiate and acquire distinct identities. A future pseudotime trajectory analysis will allow us to better understand the transition from meristematic cell to differentiated phloem or xylem cells types. Some clusters showed gradients, indicating a transition, in which transcriptional modules (for example, for xylem identity) decreased in favor of phloem modules. One example is the PC, in which cells closer to the xylem shared some xylem properties, while those closer to PP shared PP properties. A similar behavior was seen for the VP. It is conceivable that the well-studied dorsoventral cues, or cues from neighboring cells, are responsible for these gradients.

From their positioning inside the phloem, one may naively have expected the PP and CC to cluster together. CC are unique since they have to fulfil their own tasks, e.g. import sucrose from PP, and at the same time maintain the function of the adjacent enucleate sieve elements, living cells that act as conduits for assimilate translocation^56^. Based on this dual role, it is possibly not surprising that they form such a distinct island. Whether PP and CC derive from a common ancestor remains unclear since the ontogeny of PP is unknown. The ontogenesis of CC is much better understood^37^. SE/CC mother cells divide asymmetrically to produce CC and SE. The APL transcription factor is a master regulator for SE/CC development, and a repressor of xylem identity^37^. Based on our data, it seems unlikely that APL is responsible for driving PP ontogeny but may implicate alternative pathways for PP differentiation.

A major discovery was the assignment of Clusters 10 and 18 as PP, providing first insights into the PP transcriptome. The PP has essential roles in sucrose transfer to the SE/CC, but also likely many other functions in transport, metabolism and signaling. Using *SWEET11* and *12* as markers, two distinct clusters were identified, C10.2 and C18.1. It will require more careful analyses to determine if the two clusters represent spatially different cells types or developmental trajectories. *SWEET13* is a gene that may act in a compensatory role, as evidenced by higher mRNA levels in *sweet11;12* mutants. Here we detected *SWEET13* in the same cells as *SWEET11* and *12* (ref. ^38^). Notably, in a parallel study we found that while in main veins and rank-1 intermediate veins of maize, the orthologs in maize, named *ZmSWEET13a, b* and *c*, are likely also expressed in PP, in the C4 specific rank-2 intermediate veins that are mainly responsible for phloem loading, the three *ZmSWEET13*s are not in PP, but in two abaxial bundle sheath cells^57^. These data indicated to us that maize uses a different path for phloem loading of sucrose as compared to for example Arabidopsis or potato^58^. In addition to the clade III SWEETs, scRNA-seq provided extensive insights into other genes enriched in PP, in particular identifying transcripts of seven members of the UmamiT amino acid transporter family. UmamiTs are rather non-selective transporters involved at sites where cellular amino acid efflux is required. For example, *UmamiT18/SIAR1* transcripts, similar to *SWEET11* and *12*, were detected in the chalazal region of developing seed^40,59^. Knockout mutants showed reduced accumulation of amino acids in seeds, likely due to both a reduction in efflux from leaves and reduced efflux from the chalaza^40^. Of note, the *SWEET*s and *UmamiT*s showed high coexpression indices. One may thus hypothesize that the genes share transcriptional control, and thus it will be interesting to analyze whether common transcription factors binding sites are present in the PP-specific genes. SWEETs and SUTs seem to act as a pair, one responsible for sucrose efflux from PP, the other for active import into the SE/CC. UmamiTs also seem to act in pairs, with UmamiTs in PP and AAP amino acid H^+^/symporters^60^ including *AAP2, 4* and *5* in the CC (Supplementary Fig. 14). These complementary functions are consistent with the clear separation of the clusters in the UMAP plots. A striking result from the comparison of PP and CC was that their metabolism was also highly differentiated. In particular, we found major differences regarding amino acid metabolic pathways. Both isotope labeling studies as well as amino acid localization by 2D-NMR had indicated that amino acids behave very differently regarding entry into metabolism along the translocation pathway and entry into the phloem^47,48^. This difference could thus likely be explained by the metabolic activities along the path, rather than their non-selective transporters.

Cell type-specific metabolic pathway analysis revealed many other interesting aspects. For example, hormone metabolism varied between cell types. Cell types responsible for hormones biosynthesis are largely unknown. The metabolic analysis provides a map that may guide identification of hormone biosynthesis and transport pathways in leaves. For instance, high activities of ABA, ethylene, JA, and GA biosynthesis pathways were detected in C18 (which includes the PP), consistent with the high number ABA, JA, and GA transporters in the phloem^61–64^. As most transporters show functional redundancy and display diverse substrate specificity, this dataset serves as a source to identify cell type-specific redundant family members for the generation of multiple mutants, to eliminate or verify interacting partners, and to identify yet -unknown transporters.

Besides the insights into both metabolism and into apoplasmic transport pathways, the study also showed that cell types with unique types of plasmodesmata express particular paralogs of plasmodesmata-specific proteins, such as PDLPs or MCTPs. The combined analysis of sym-and apoplasmic fluxes of ions, metabolites and signaling molecules at the cellular level will likely enable a much better understanding of the physiology of the leaf.

In summary, scRNA-seq enabled us to identify unique and distinct features of the different vascular cell types present the leaf. Through transcriptomic and metabolic pathway analyses at the single cell level, we identified potential roles of PP not only in the transport of sugars, but also in amino acid transport. Importantly, we also identified unexpected roles of PP in hormone biosynthesis and defense-related responses. The information provided in this study provides key resources to develop strategies influencing the flux of ions, metabolites and signals.

## Methods

### Enrichment of vascular protoplasts

Protoplasts were isolated from mature leaves of 6-weeks-old Arabidopsis Col-0 plants grown under short-day (8 h light / 16 h dark) conditions at a PAR of 60 µmol m^-2^ s^-1^. For protoplast isolation, the tape-sandwich method^15^ was modified and applied for removal of the abaxial epidermis, guard cells and trichomes. The adaxial side of the fully developed leaves was stabilized by placing on the time tape, and the abaxial side was adhered to the 3M Magic tape. The abaxial epidermis was removed by pulling off the 3 M Magic tape. Two cuts were made on each side of the major vein of the peeled leaves using a razor blade (Swann-Morton, Sheffield). Seven to nine abaxial epidermis-peeled and cut leaves attached to time tape were immediately immersed in the petri dish containing 15 mL freshly prepared protoplast isolation solution (1% Cellulase Onuzuka R-10 (Duchefa, Haarlem), 0.3% macerozyme R-10, (Duchefa, Haarlem), 0.6 M Mannitol, 20 mM MES pH 5.7, 20 mM KCL, 1 mM DTT, 10 mM CaCl_2_, and 0.1 % BSA). Dithiothreitol (DTT) and bovine serum albumin (BSA) were added to the enzyme solution to protect the protoplasts. Leaves were shaken at 30 rpm on a platform shaker for 2 hours. The release of the protoplasts was monitored every 30 min by checking released cells under the microscope or by monitoring what was left on the time tape. After confirming the release of the protoplasts into the solution, 10 mL of wash buffer (0.6 M Mannitol, 20 mM MES pH 5.7, 20 mM KCl, 1 mM DTT, 10 mM CaCl_2_, and 0.1 % BSA) was slowly added to the petri dish containing the protoplasts. The solution was filtered into a round bottom 20 mL tube using a 70 μm pore size filter (Corning, New York). The protoplast solution was centrifuged at 100 x *g* for 3 min in a swinging rotor. The protoplasts were washed four times with 20 mL washing buffer. After the final wash step, protoplasts were slowly resuspended in 1 mL wash buffer and gently filtered twice using a 40 μm Flowmi cell strainer (Bel-Art SP Scienceware, New Jersey). The number of protoplasts was counted with the C-Chip Neubauer improved hemocytometer (NanoEnTek, Seoul) under a light microscope (Leica VT1000, Wetzlar). The viability of protoplasts was determined using trypan blue solution (Gibco; Thermo Fisher Scientific, Massachusetts).

### Quantitative reverse transcription (RT-qPCR)

Total RNA was extracted using the RNeasy Kit (Qiagen, Hilden). cDNA synthesis and gDNA removal steps were performed using QuantiTect Reverse Transcription Kit (Qiagen, Hilden), qPCR was performed on Stratagene Mx3000P (Agilent Technologies, California) using the Lightcycler 480 SYBR Green I Master Mix (Roche, Penzberg). Transcript levels were quantified using the relative standard curve method. Values were normalized to the level of the internal control, *UBQ10*. Primers used for RT-qPCR are listed in Supplementary Table 15. For RT-qPCR of *apl* mutants, segregating seeds from heterozygous parents were sown on MS media. RNA was extracted using 2-weeks-old plants grown under LD conditions.

### Single cell RNA-seq library preparation and sequencing

Two biological replicates were performed with the aim of capturing ∼7,000 leaf protoplasts for each replicate. Freshly isolated protoplasts were adjusted to 700-900 cells/µL and loaded into the 10X Genomics Chromium single cell microfluidics device according to the Single Cell 3’ Reagent Kit v2 protocol (10x Genomics, California). Eleven cycles were used for cDNA amplification and 12 cycles were used for final PCR amplification of the adapter-ligated libraries. The quality and size of the final library was verified on a DNA High Sensitivity Bioanalyzer Chip (Agilent Technologies, California), and libraries were quantified using the NEBNext Library Quantification Kit for Illumina (New England Biolabs, Massachusetts). scRNA-seq library sequencing was performed on a NextSeq platform (Illumina Inc, California), using the sequencing parameters 26,8,0,98 (c.ATG, Tübingen).

### Generation of single cell expression matrices

Reads were aligned to the *Arabidopsis thaliana* reference genome (Araport 11) using Cell Ranger 3.0.2 (10X Genomics, California) with default parameters. The output files for the two replicates were aggregated into one gene-cell expression matrix using Cell Ranger aggregate with the mapped read depth normalization option.

### Dimensionality reduction, UMAP visualization, cell clustering analysis, and correlation analysis

The Seurat R package (version 3.1.0)^17,65^ was used for dimensionality reduction analysis. The SCTransform option was used for normalization, scaling the data and finding variable genes using default parameters^66^. During normalization, potential variation due to mitochondrial mapping percentage was removed. We did not remove rRNA before sample preparation and did not filter cells with higher rRNA transcript levels, since rRNA levels can vary among cells. Fifty principal components (PCs) were selected as input for a graph-based approach to cluster cells by cell type using a resolution value of 0.8 in all clustering analyses. Uniform Manifold Approximation and Projection (UMAP) dimensional reduction^18^ was used for two-dimensional visualization using ten PCs, 30 neighboring points and a minimum distance of 0.1. Subclustering was performed using the same parameters. For the correlation analysis between single cell replicates across individual clusters, the average expression of cells within a cluster was calculated and the Pearson-correlation coefficient was determined.

### Identification of Differentially Expressed Genes and Cluster-Specific Marker Genes

Genes differentially expressed across clusters or subclusters were identified by comparing average transcript levels in cells of a given cluster to that of cells in all other clusters using the Seurat package likelihood ratio test (Bimod). The following cutoffs were applied: average expression difference ≥0.25 natural Log and q <0.01. Cluster-specific marker genes were selected from among the differentially expressed genes based on the criteria that marker genes must be expressed in >10% of cells within the cluster (PCT1), and <10% of cells across all other clusters (PCT2).

### Bulk RNA-seq library preparation and sequencing

Total RNA was extracted from leaves (not protoplasted) and leaf protoplasts isolated using the same method for the single cell sequencing using the RNeasy Kit (Qiagen, Hilden). On column DNase treatment was performed to remove residual gDNA using the RNase-free DNase kit (Qiagen, Hilden) following manufacturer recommendations. Two biological replicates were made for leaf and leaf protoplast samples. The integrity of the RNA was confirmed using Agilent RNA 6000 Nano Chip (Agilent Technologies, California) and LabChip GX (PerkinElmer, Massachusetts). RNA concentration was measured using the Qubit Fluorometer using the RNA broad range quantification kit (Thermo Fisher, Massachusetts). For mRNA poly-A enrichment, 5 μg of total RNA was purified using the Poly(A) mRNA Magnetic Isolation Module (New England Biolabs, Massachusetts). Libraries were constructed using the ULTRA II directional library kit (New England Biolab, Massachusetts), and size selection was done using SPRI beads (New England Biolab, Massachusetts) following the manufacturer manual with the following exceptions: 7 min of fragmentation time and 26.5 and 10 μL SPRI beads pre-PCR were used to enrich ∼400 bp inserts. Library amplification included 10 PCR cycles, and 0.7 volumes of (35 μL) of SPRI beads were used for post-PCR purification. QC-tested libraries (Agilent Technologies, California) were sequenced on an Illumina HiSeq 2500 lane with 150 bp paired-end (Novogene, Beijing).

### Bulk RNA-seq analysis

Paired-end reads (150 bp) were aligned to the *Arabidopsis thaliana* reference genome (Araport11) using STAR^67^ (maximum intron length of 2 kbp). Differential expression analysis was carried out in R (v3.6.1) using Bioconductor (v3.9) and DESeq2 (v1.24) (absolute Log2FC ≥ 1 and q value < 0.05). For correlation analysis of gene expression between protoplasted and non protoplasted bulk tissues, the Log2 (mean FPKM+1) expression values were calculated for each gene. Pearson-correlation coefficient was determined in R.

### Pathway activities in different cell types

We followed the formulation introduced in Xiao *et al*.^49^ to calculate a pathway activity score, which depends on the mRNA level of its constituent genes. The mRNA count of gene *i* in cell *k* was denoted as *g*_*i,k*_. We first normalized the mRNA count of a gene in a cell by the average for all genes in the cell, and denoted this normalized level as *g’*_*i,k*_. The mean transcript level *E*_*i,j*_ of gene *i* in cell type *j* is then defined as the mean *g’*_*i,k*_ over all cells of that type, 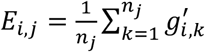, where *n*_*j*_ is the number of cells classified as cell type *j*, and *k* is the index for individual cells. *E*_*i,j*_ is normalized by the average transcript level of gene *i* across all cell types to become the relative transcript level *r*_*i,j*_ of gene *i* in cell type 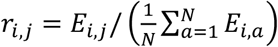, where *N* is the total number of cell types. The pathway activity score (PAS) of pathway *t* in cell type *j*, denoted as *p*_*t,j*_, is then a weighted average of the relative transcript levels across the pathway genes: 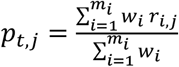; here, *m*_*t*_ is the number of genes in pathway *t*, and the weight *w*_*i*_ of gene *i* is defined as the reciprocal of the number of pathways that include gene *i*, to ensure a stronger influence of pathway-specific genes. As the relative mRNA levels *r*_*i,j*_ are centered around 1, the same is true for the pathway activity *p*_*t,j*_, with *p*_*t,j*_ <1 corresponding to underrepresentation of pathway *t* in cell type *j* relative to the pathway’s activity across all cell types; conversely, *p*_*t,j*_ >1 indicates a higher than average activity in cell type *j*. To assess the statistical significance of a *p*_*t,j*_ value, we performed a permutation test, shuffling the cell type labels of the genes a thousand times to simulate the null distribution of *p*_*t,j*_ under the assumption of no systematic cell type specific pathway activity; we defined an empirical *p*-value by comparing *p*_*t,j*_ to this null distribution. AraCyc pathway lists^68^ used for the analysis can be found in Supplementary Table 14.

### Generation of reporter lines

For the generation of *pbZIP9:GFP-GUS* lines, a 2,289 bp fragment of the *bZIP9* gene promoter was amplified using bZIP9pro attB1 and bZIP9pro attB2 primers using Col-0 genomic DNA as template. The corresponding PCR fragment was purified and used for GATEWAY BP reaction into pDONR221 and cloned into the destination vector pBGWFS7,0 through LR reaction. For generating *pSWEET11:SWEET11-2A-GFP-GUS* lines, a 4,784 fragment consisting of promoter and SWEET11 genomic region including all exons and introns was amplified using SWEET11-2A-attB1 and SWEET11-2A-attB2 primers. The SWEET11-2A-attB2 reverse primer contained the 2A cleavage sequence. The genomic SWEET11 with the 2A cleavage site was cloned into pDONR221 and subcloned into the pBGWFS7,0 vector through LR reaction. To generate *pSWEET13:SWEET13-YFP*, a 3,941 bp fragment including SWEET13 promoter and its genomic region was amplified using SWEET13attB1 and SWEET13attB2 primers and cloned into the donor vector pDONR221-f1 by BP reaction. Subsequently, the fragment was sub-cloned into a gateway-compatible vector, pEG-TW1^69^ to generate pSWEET13:SWEET13-YPF construct by LR reaction. Transformation of plants was performed using the floral dip method^70^. Primers used for amplification are listed in Supplementary Table 15.

### GUS histochemistry

GUS staining was performed as described with minor modifications^71^. Tissues were fixed with 90% ice-cold acetone. After applying vacuum for 10 min, acetone was removed and replaced by prestaining solution (1 mM EDTA, 5 mM potassium ferricyanide, 5 mM potassium ferrocyanide, 100 mM sodium phosphate (pH 7.0), 1% Triton-X-100). After 10 min vacuum, the prestaining solution was replaced with staining solution containing 1 mM EDTA, 5 mM potassium ferricyanide, 5 mM potassium ferrocyanide, 100 mM sodium phosphate (pH 7.0), 1% Triton-X-100, and 2 mM X-Gluc and further vacuum infiltrated for 15 min in the dark. The tissues in staining solutions containing X-Gluc were incubated at 37 °C in the dark for 4 hours. Ethanol series were performed from 25% to 70% ethanol in 30 min steps.

### Sample preparation for imaging leaf vasculature

Rosette leaves were excised by cutting the leaf petiole from 5 weeks old plants grown under SD conditions. Double-sided tape was used to attach the petiole and the upper lamina of the leaf on the slide glass (abaxial side facing upward, as in the tape-sandwich method). The abaxial epidermis was peeled using the 3M Magic tape and immediately covered with water. The region of interest was excised with a razor blade and moved to a new slide glass. Samples were applied with FM4-64FX (Sigma-Aldrich, Missouri) (5 μg/ml) for 5 min for staining the cell membrane and imaged immediately. PP-cells were identified based on the cell size and chloroplast organization as described^8^.

### Confocal imaging

Fluorescence images were captured using a Leica TCS SP8 confocal microscope with a 20x or 40x objective with water immersion. GFP, YFP, FM4-64FX, and chlorophyll autofluorescence signals were acquired using the following settings: GFP, excitation 488 nm (white-light laser) and emission 492-552 nm; YFP, excitation 514 nm and emission 510-565 nm; FM4-64FX, excitation 561 nm and emission 599– 680 nm; chlorophyll autofluorescence, excitation 638 nm and emission 645-738 nm.

### Reporting Summary

Further information on research design is available in the Nature Research Reporting Summary linked to this article.

## Data availability

The raw data that support the findings of this study are available from the corresponding author upon reasonable request. All sequencing data have been deposited in the Gene Expression Omnibus GEO (www.ncbi.nlm.nih.gov/geo/) under the accession number GEO X and The Single Cell Expression Atlas at EMBL-EBI (www.ebi.ac.ik/gxa/sc/home).

## Acknowledgements

We thank Colin P. S. Kruse (Los Alamos National Laboratory, USA) for constructive comments on the metabolic pathway activity analysis, Thomas Hartwig (WBF lab) for cDNA library preparation for bulk RNA-sequencing, Sebastian Hänsch for assistance with confocal microscope imaging (Center for Advanced Imaging, HHU, Germany), and Xiaoqing Qu (Bio-Protocol, China) for helping with the generation of pSWEET13:SWEET13-YFP construct. This research was supported by the National Science Foundation (SECRETome Project: Systematic Evaluation of CellulaR ExporT from plant cells, IOS-1546879), Deutsche Forschungsgemeinschaft (DFG, German Research Foundation) under Germany’s Excellence Strategy – EXC-2048/1 – project ID 390686111 and SFB 1208 – Project-ID 267205415, as well as the Alexander von Humboldt Professorship to WBF.

## Author contributions

J.Y.K. and W.B.F. conceived and supervised the projects. J.Y.K. and T.D. generated the scRNA-seq libraries.

E.S. and J.Y.K. performed bioinformatics analyses. J.Y.K., M.B., M.M., M.M.W. and W.B.F. analyzed clusters, J.Y.K., D.W., N.Z., and L.Q.C. generated and analyzed reporter lines. T.Y.P. performed metabolic pathway activity analyses in consultation with M.J.L., J.Y.K. and W.B.F. wrote the manuscript. All authors discussed the results and commented on the manuscript.

## Supplementary information

Supplementary Text

Supplementary Figs 1-13, Supplementary Tables 1-15, Supplementary Video 1

**Correspondence and requests for materials** should be addressed to Ji-Yun Kim.

## Supplementary text and data

### Possible limitations of the scRNA-seq

Despite the advances in scRNA-seq technologies, application to plant cells still faces challenges since the cell walls of plant cells must be removed for the release of individual cells and the penetration of the buffers into cells. Different cell types are likely under different osmotic pressure in the intact plant which might result in breakage if the osmotic conditions are not adequate. Recent studies using Arabidopsis^73^, rice^74^, as well as maize^57^ aerial tissues clearly demonstrate difficulty in capturing vascular cell types reflecting the need for careful strategy development. An unexpected challenge we faced while using loading the cells to the microfluidic chip (10X Genomics) was the sedimentation of mesophyll cells (size of < 50 um). This could also be true for cells from other species but not evident due to the transparent or light color of the cells. However, due to the sedimentation of cells, we observed a 30% loss in the recovery in one of our replicates (Replicate 2). Both libraries showed a high correlation and were therefore used for further analysis. The development of plant cell-specific strategies and compatible buffers will reduce the failure rate and facilitate the application of scRNA-seq analysis in plant tissues from diverse species.

### Transcripts enriched in the procambial cluster

Several factors known to be expressed in the phloem precursor cells and play roles in the acquisition of phloem identity such as *ALTERED PHLOEM DEVELOPMENT* (*APL*)^37^, a transcription factor known to promote phloem differentiation, and phosphoinositide 5-phosphatases *COTYLEDON VASCULAR PATTERN 2* (*CVP2*) and *CVP2 LIKE 1* (*CVL1*)^34^ were detected in the central region of Cluster 10 (C10.1.1) (Fig. 4, Supplementary Fig. 5). *BRX* and *OCTOPUS*^75^, important factors involved in protophloem differentiation expressed at different ends of the developing protophloem cells, were also enriched in the nonoverlapping cells in C10.1.1. The protophloem specific *CLAVATA3/EMBRYO SURROUNDING REGION-RELATED 45* (*CLE45*) peptide required for the maintenance of pluripotent phloem cell reservoirs through the interaction with the leucine-rich repeat receptor kinase (LRR-RK), *BARELY ANY MERISTEM 3* (*BAM3*)^35,36^, was also detected in subset cells of the procambium cluster. The *CLE45*-enriched cells are in the lower file within the central region (C10.1.1), whereas the *BAM3*-enriched cells are broadly distributed in Cluster 10 with the highest expression in cluster C10.1.1. The distribution of *CLE45* and *BAM3*-expressing cells in our UMAP is reminiscent of the overlapping and nonoverlapping distribution pattern of *CLE45* and *BAM3* in roots^76^ suggesting that the mechanism for retaining the plastic identity of the root might also be present in the leaf.

Proper control of cell division and proliferation rate is a critical factor for maintaining the meristematic state of the procambium. The auxin response factor MONOPTEROS (MP)^77^ plays a role in vascular proliferation by activating of *TARGET OF MONOPTEROS 5* and the auxin and cytokinin responsive DOF family transcription factor *TARGET OF MONOPTEROS 6 (TMO6). TMO6* is further regulated by the receptor kinase PHLOEM INTERCALATED WITH XYLEM and is part of the feed-forward loop involving WUSCHEL HOMEOBOX RELATED 14 (WOX14), TMO6 and LATERAL ORGAN BOUNDARIES DOMAIN 4 (LBD4). This feed-forward loop takes place the boundary of the phloem and procambium cells and influences the distribution of phloem by promoting cell division ^78^. We detected *WOX14* transcripts in a few cells at the central region of the procambium cluster. *LBD4* and *TMO6* transcript are present in the boundary of phloem (PP, C10.2) and procambium cell cluster (C10.1.1). Others, including the plant aurora kinases that affect phloem and xylem differentiation negatively through cell division rate control^79^, are both enriched in the procambium cell cluster in C10.2.1(Supplementary Fig. 5). Together, we demonstrate that the leaf single-cell sequencing analysis reveals different cell identities within the procambium cluster expressing marker genes identified in various tissues (roots and stems) and show that the spatial distribution of the clusters is well correlated with the distribution of the cell types *in planta*. UMAP plots of additional procambium marker genes in the subpopulations of C10 are shown in Supplementary Fig. 5.

### Cluster 19 as a candidate for putative S-cell cluster

Glucosinolate-rich cell type (S-cells) are found in floral stems of Arabidopsis but also described as sulfur-rich cells in the phloem parenchyma of the leaf ^50^. S cells contain more than tenfold higher sulfur levels, and high levels of glucosinolate compared to the rest of the cells and have been the functional shield of the plant vascular system from herbivores and pathogens. As the S-cells were suggested to be found in the abaxial area of phloem parenchyma in leaves, we questioned whether the S-cells are present in the PP clusters. Markers for S-cells are presently unavailable, therefore, we searched for markers that are known to be absent in S-cells. Proteomic analysis of S-cells has suggested that glucosinolates are not produced in S-cells, as biosynthetic enzymes could not be detected in isolated S-cell extracts. The promoter activities of key enzymes and markers for glucosinolate biosynthesis such as *CYTOCHROME P450 83A1*(*CYP83A1*) and *BRANCHED-CHAIN AMINOTRANSFERASE 4* (*BCAT4*) were not detected in S-cells^81–83^. We detected transcripts of *CYP83A1, BCAT4*, as well as other transcripts encoding proteins involved in the synthesis of aliphatic and indolic glucosinolates in the bundle sheath, xylem (C4) and PP/PC/XP (C10, C18) clusters (Supplementary Fig 12). The enrichment of *HIG1* transcription factor which activate promoters of genes involved in glucosinolate biosynthetic genes^84^ (Supplementary Tables 4 and 7) also exclude the cells in the PP clusters (C10.2, C18.1) from containing S-cells and suggest the phloem parenchyma, procambium, xylem, and bundle sheath cells as sites for glucosinolate biosynthesis.

A unique characteristic of the S-cells is that these cells undergo programmed cell death at the early stages of differentiation. We therefore, searched for cells that could be enriched with the transcripts related to programmed cell death. We identified Cluster 19, a cluster distinct from, but closely spaced to the bundle sheath cells. This cluster was enriched with transcripts related to programmed cell death, hypersensitive response, and defense and immune response (Supplementary Table 13). This cluster also showed high activity scores in insect chewing-induced glucosinolate breakdown pathway (Supplementary Fig. 9b). This result is in line with the primary role of S-cells in releasing toxic compounds through glucosinolate breakdown upon chewing insects induced-mechanical disruption. Although we cannot rule out that these subset cells were clustered based on the stress response from the protoplasting process, they could serve as candidates as putative S-cells.

### Comprehensive PAS analysis across all cell types

A complete PAS analysis is shown in Supplementary Figure 10.

### Highlights from the PAS analysis

- Glucosinolate biosynthesis activity is high in Cluster 10, Cluster 18, and Cluster 4.
- The pathway activity score of several glucosinolate biosynthesis pathways are low in the GC and CC.
- Mesophyll cells are unlikely sites for glucosinolate biosynthesis nor breakdown.
- The pathway activity scores of glucosinolate breakdown pathways are high in the guard cells.
- Cluster 19 is enriched in glucosinolate breakdown pathway induced by insect chewing.
- Cluster 18 (PP2 and XP3) is enriched with ABA, Ethylene, JA, GA biosynthesis pathways.
- Clusters 10 and 18 are enriched with callose biosynthetic pathways.
- The activity scores of photosynthesis-related pathways are low in non-mesophyll cells (epidermis, CC, guard cell, PP, PC, XP3).

---

**Supplementary Fig. 1:**
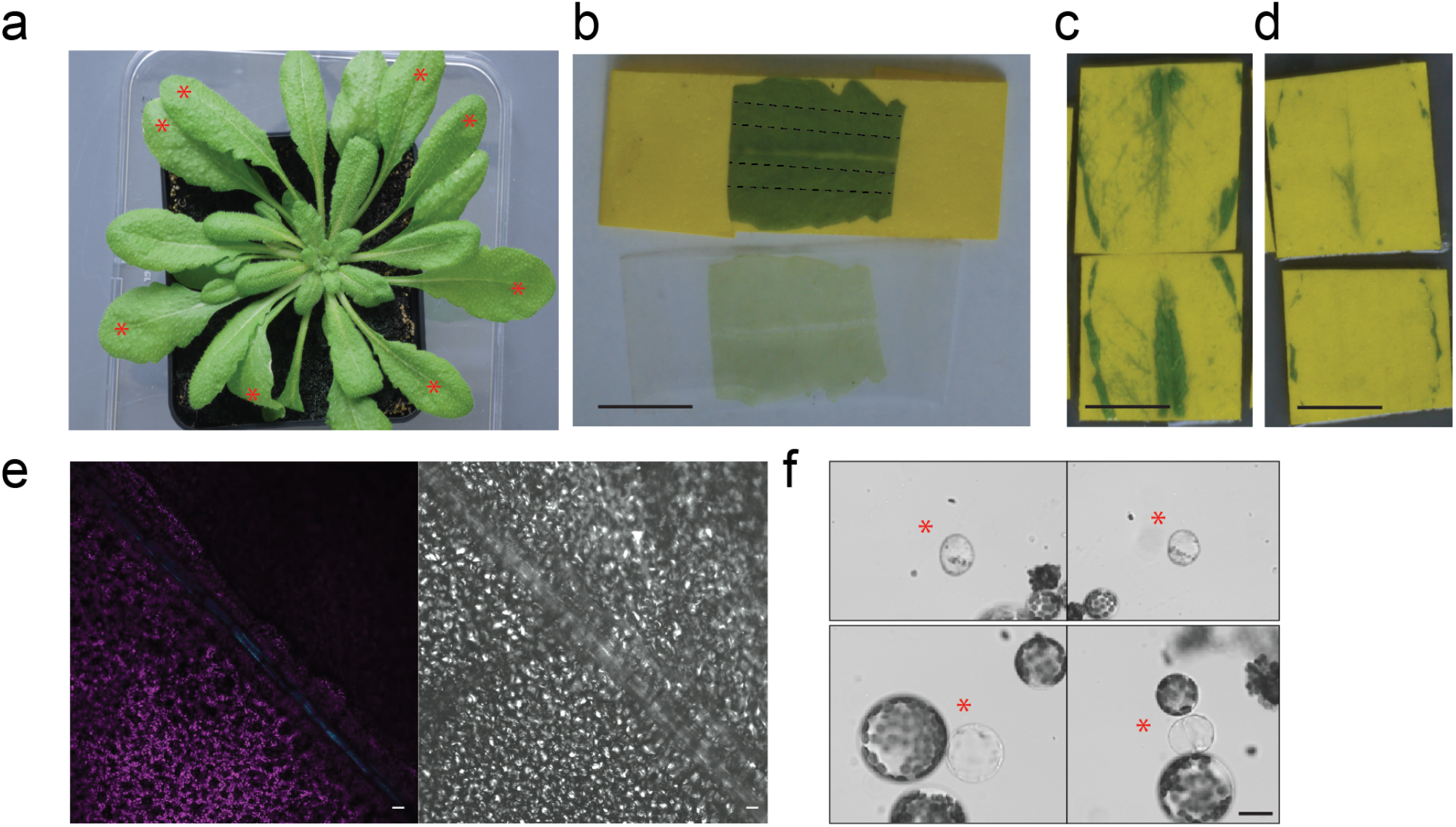
Optimization of vascular protoplast enrichment. **a**, Six-week-old Arabidopsis plant grown under short-day conditions. Leaves used for protoplasting are marked with red asterisks. **b**, Tape-sandwiched leaf explants were separated and the remaining adaxial side was cut parallel to the main vein. Cutting sites are marked as dotted lines. Scale bar: 1 cm. **c-d**, Tape-separated leaf explants 2 hours after enzyme digest. The release of vascular cells is more effective with an enzyme solution in 0.6 M mannitol (**d**), compared to 0.4 M mannitol (**c**) Scale bar: 1 cm. **e**, Detection of procambial cells in the major vein using the procambial marker *Q0990*. Scale bars: 50 μm. **f**, Diversity of leaf protoplasts. Protoplasts lacking chloroplasts are marked with red asterisks. Scale bar: 20 μm

**Supplementary Fig. 2:**
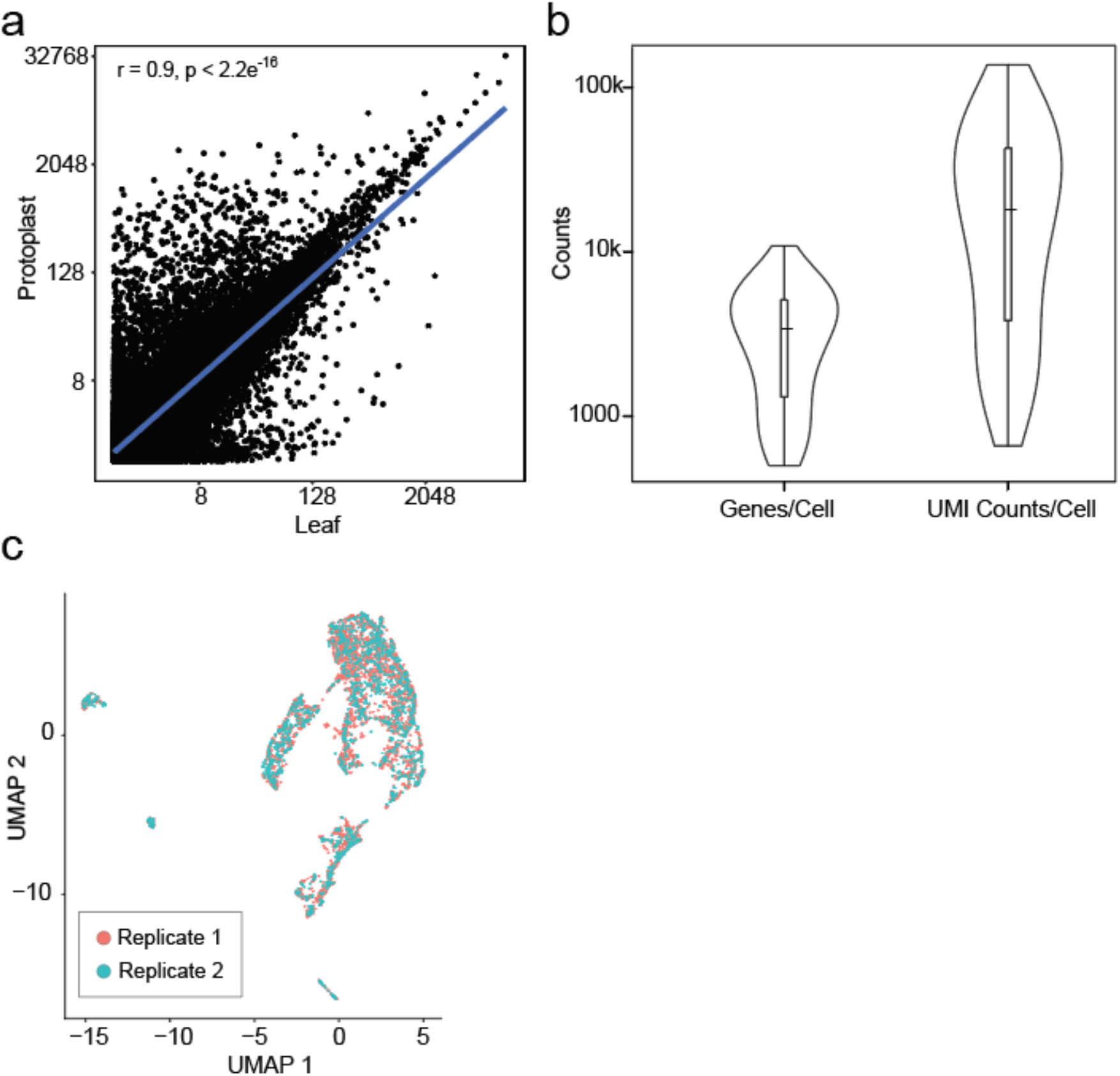
Reliability of Arabidopsis leaf scRNA-seq dataset. **a**, Correlation of gene expression in bulk leaf (not protoplasted) and bulk protoplasts. r = Pearson’s correlation coefficient. The list of DEGs in bulk RNA-seq of non protoplasted leaf and protoplasted leaf samples are provided in Supplementary Table 1. **b**, Violin plot illustrating the distribution of the number of genes (nFeature RNA) and UMI counts (nCount RNA) detected within a cell. **c**, UMAP plot of 5,230 cell transcriptomes from two biological replicates. Different colors indicate the cells from each replicate.

**Supplementary Fig. 3:**
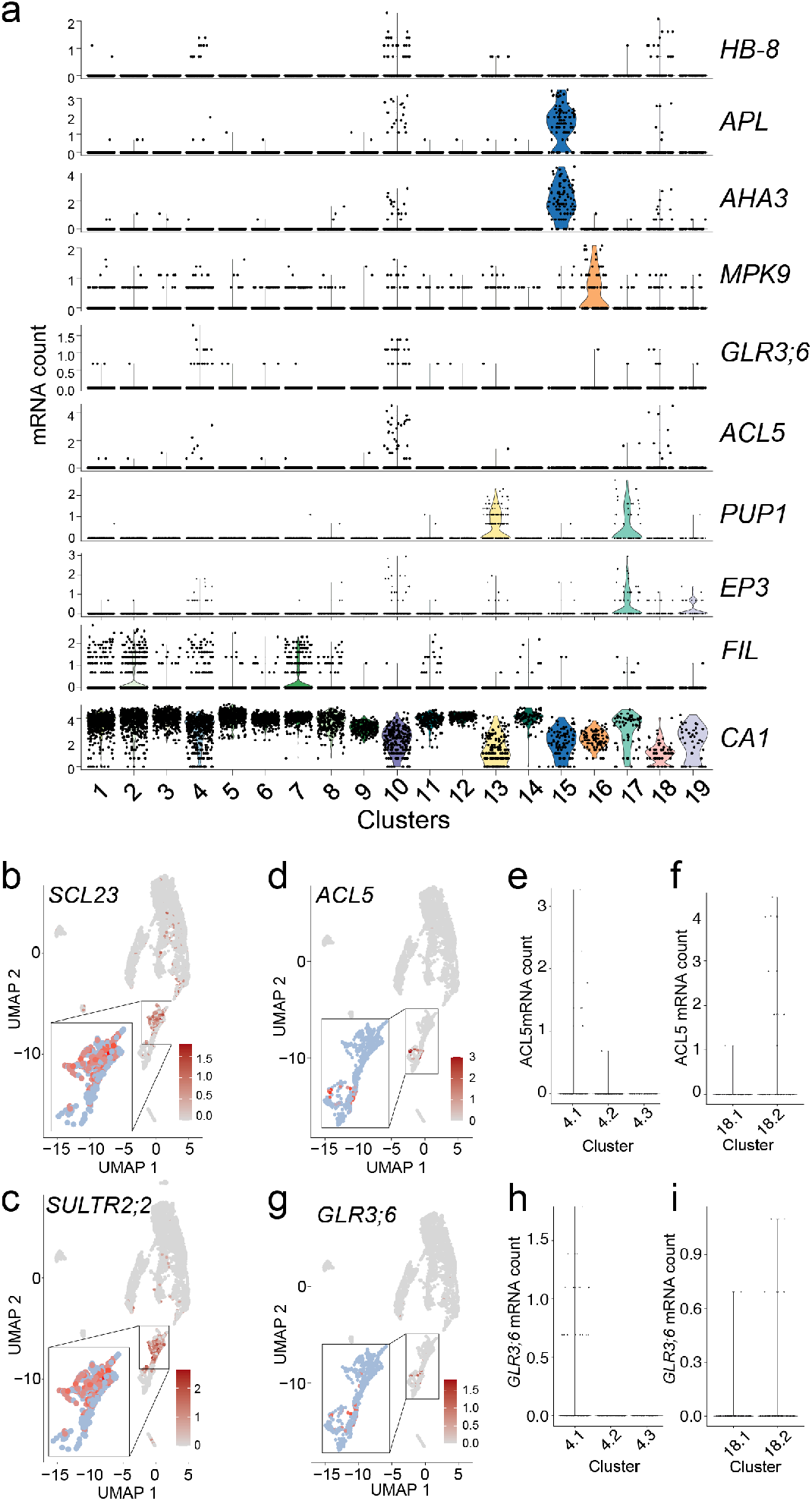
Enrichment of known marker genes for assigning cell types to clusters. **a**, Violin plots showing the expression of cell type-specific marker genes across clusters. **b**,**c**, UMAP plot illustrating the enrichment of bundle sheath (BS) markers, *SCL23* (**b**) and *SULTR2;2* (**c**). Inset shows magnification of Cluster 4. **d-i**, UMAP plot (**d**,**g**) and violin plots of the subclusters of Cluster 4 (**e**,**h**) and Cluster 18 (**f**,**i**) showing enrichment of xylem markers, *ACL5* (**d**,**e**,**f**) and *GLR3;6* (**g**,**h**,**i**). Magnified views of the boxed region (Clusters 4, 10, and 18) is shown in the insets (**d**,**g**).

**Supplementary Fig. 4:**
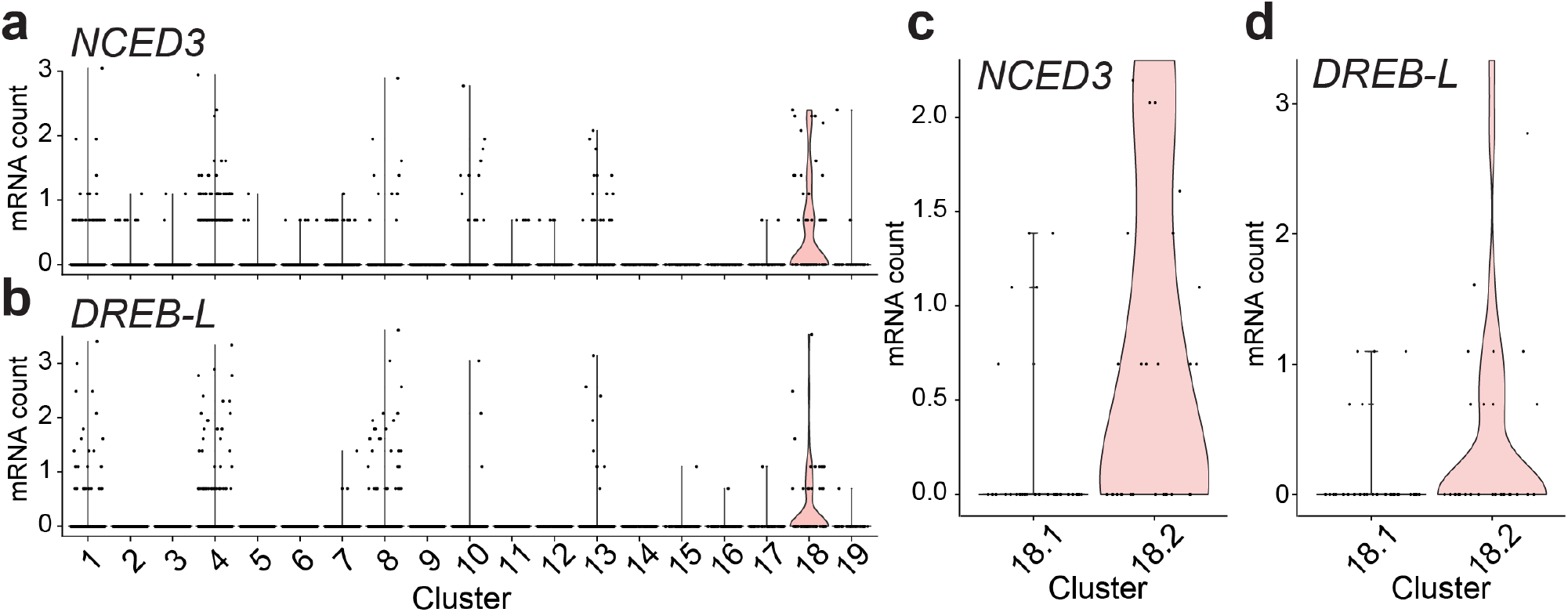
Vascular parenchyma-specific marker genes enriched in Cluster 18 (XP3) **a-d**, Violin plots showing the transcript enrichment of *NCED3* (**a**,**c**) and *DREB-L* (**b**,**d**) across the main clusters (**a**,**b**) or subclusters of Cluster 18 (**c**,**d**).

**Supplementary Fig. 5:**
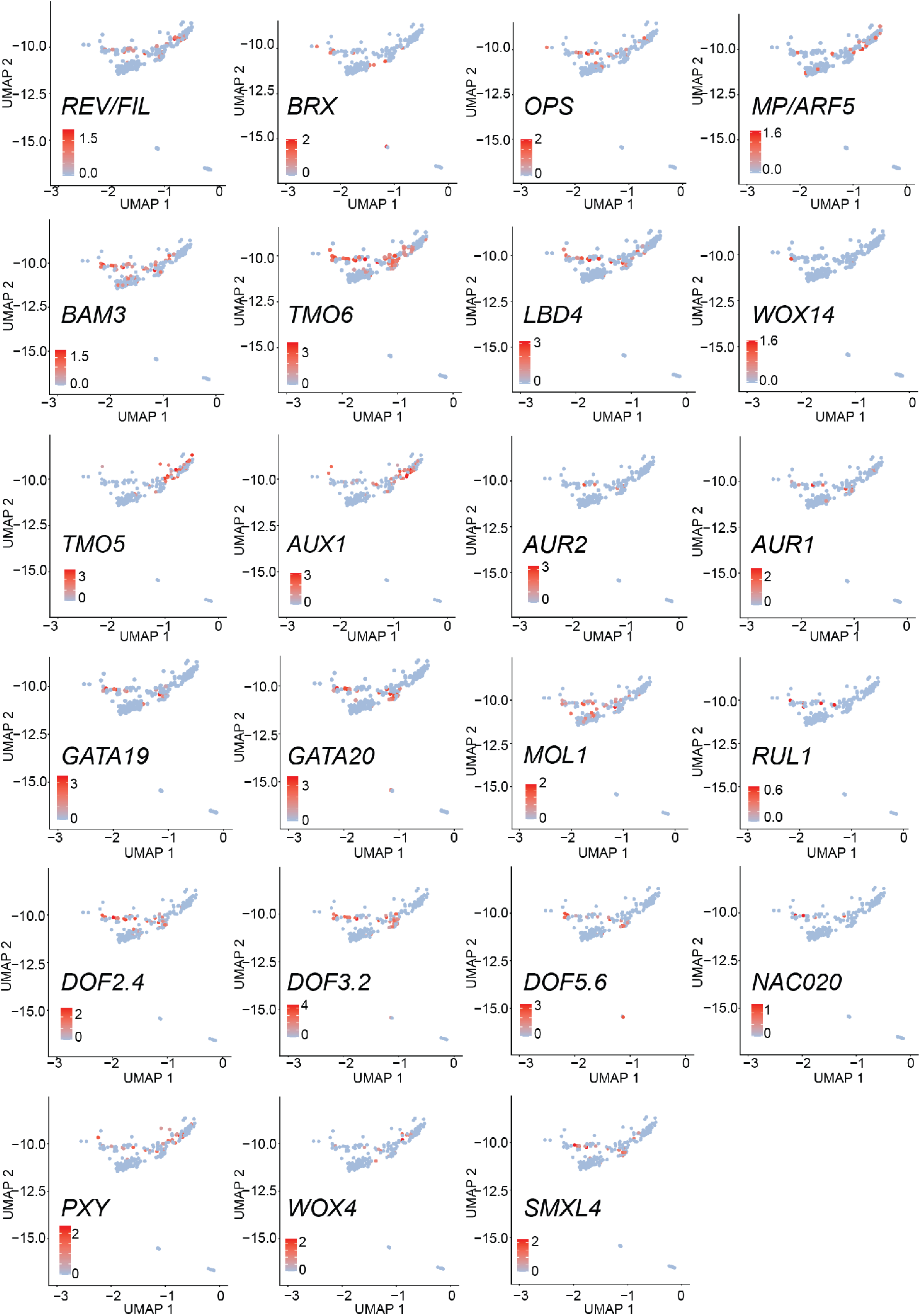
Distinct procambium cell identities in C10 subclusters. UMAP plot showing the distribution of transcripts enriched in C10 subclusters.

**Supplementary Fig. 6:**
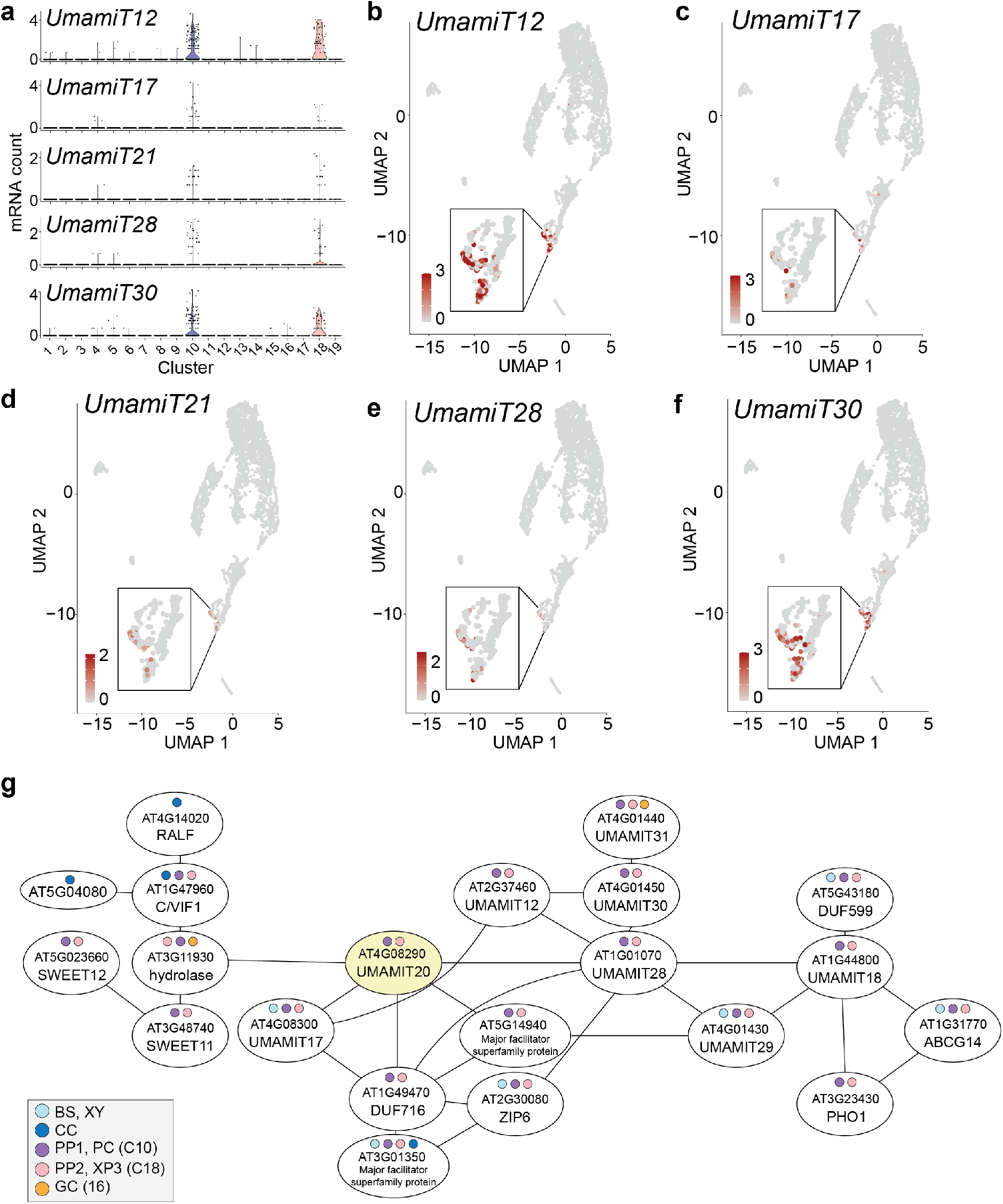
Transcripts of UmamiT amino acid transport family members are enriched in the PP. **a**, Violin plots illustrating the transcript enrichment of *UmamiT12, UmamiT17, UmamiT21, UmamiT28*, and *UmamiT30*. The cell types assigned to clusters are indicated in **Fig. 2a**,**b. b**, UMAP showing the enrichment of *UmamiT12* **(b)**, *UmamiT17* **(c)**, *UmamiT21* **(d)**, *UmamiT28* **(e)**, and *UmamiT30* **(f)**. Magnifications of the marked regions are shown in the insets. **g**, Coexpression gene network around *UmamiT20* built from the coexpression database ATTED-II (http://atted.jp). The cell types in which the genes in the network are enriched are marked with colored circles. The colors corresponding to the cell types and clusters are indicated in the bottom left box.

**Supplementary Fig. 7:**
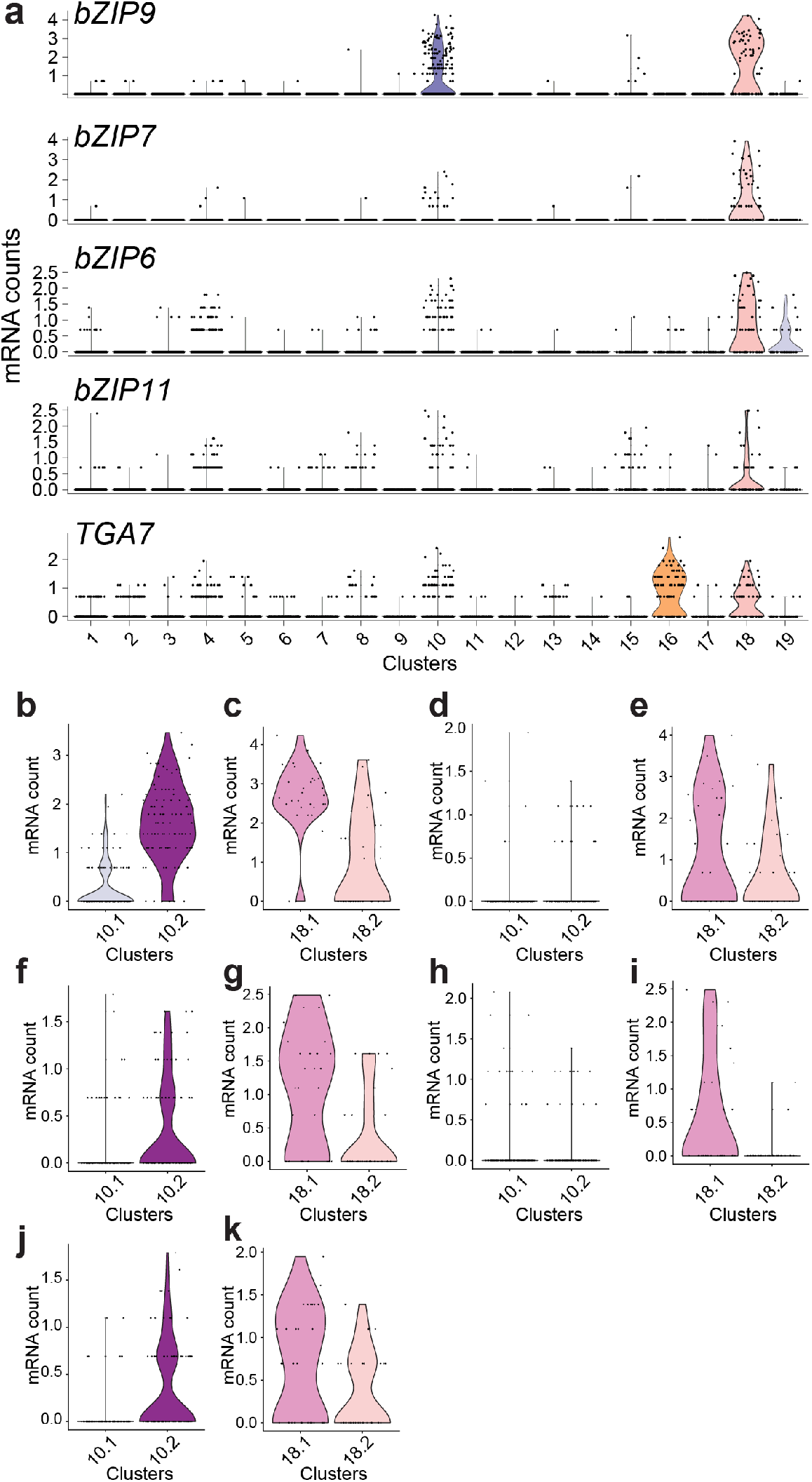
Transcript enrichment of multiple bZIP transcription factors in the PP clusters. **a**, Violin plots showing transcript enrichment of five bZIP family members. **b-k**, Violin plots showing transcript enrichment of *bZIP9* (**b**,**c**), *bZIP7* (**d**,**e**), *bZIP6* (**f**,**g**), *bZIP11* (**h**,**i**), and *TGA7* (**j**,**k**) in the subclusters of Cluster 10 (**b**,**d**,**f**,**h**,**j**) and Cluster 18 (**c**,**e**,**g**,**i**,**k**).

**Supplementary Fig. 8:**
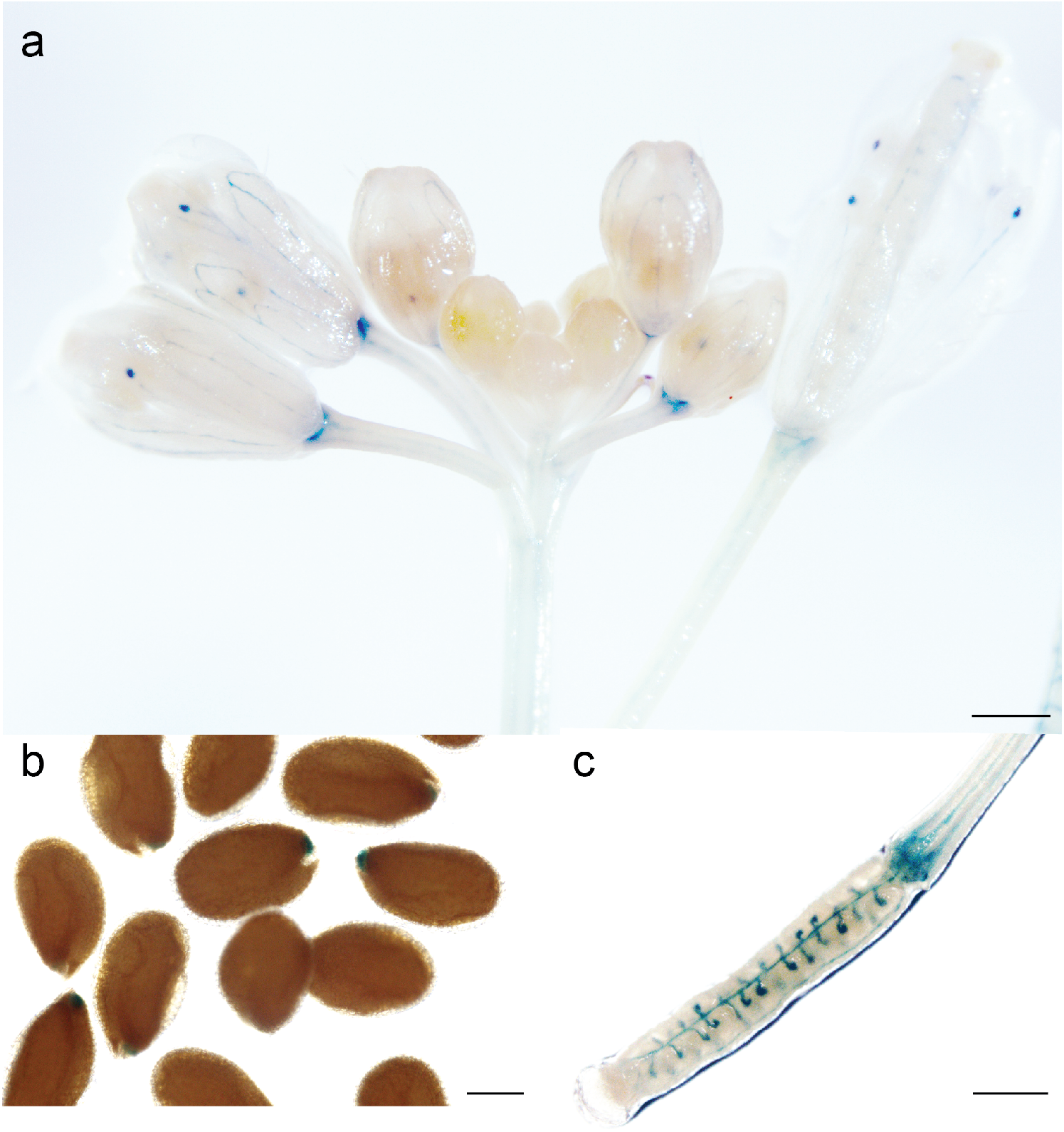
GUS staining of *pbZIP9:GFP-GUS* plants showed promoter activity in multiple tissues and organs. **a-c**, GUS stained *pbZIP9:GFP-GUS* transgenic plants show GUS activity in the unloading zone of the anther, petals, and receptacle (**a**), the unloading zone of the seed coat (**b)**, and the ovules, transmitting tract, and the funiculus of the ovary (**c**). Scale bars: 500 μm (**a**,**c**) and 200 μm (**b**).

**Supplementary Fig. 9:**
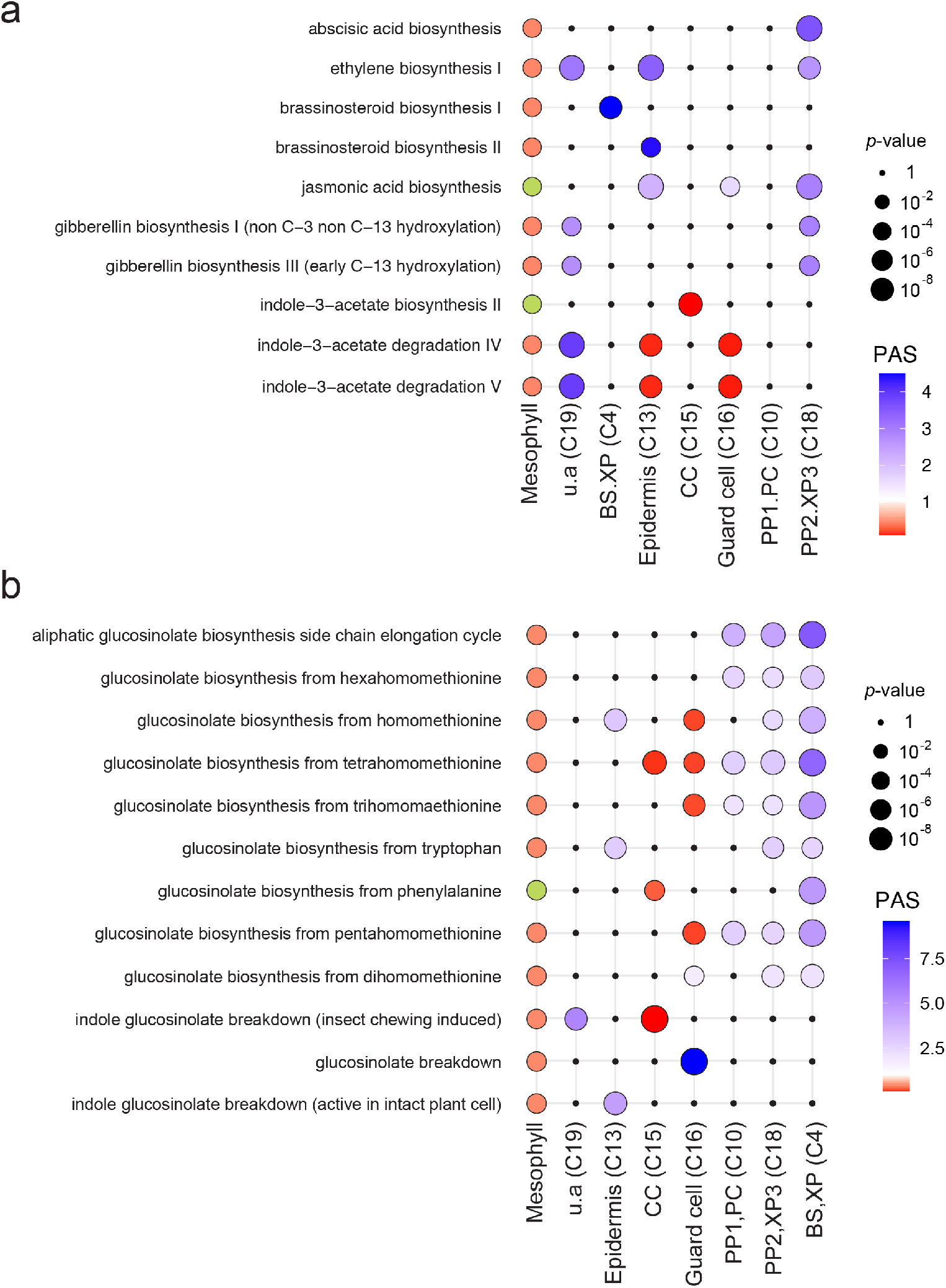
Hormone and glucosinolate pathway activity across clusters. **a**, Pathway activity score (PAS) of hormone biosynthesis and degradation pathways in the indicated clusters. **b**, Metabolic pathway activities of glucosinolate biosynthesis and degradation pathways. **a**,**b**, The color of the dots PAS, whereas the size of the dot corresponds to its statistical significance; PASs that are statistically insignificant, *i*.*e. p* >0.05 in a random permutation test, are represented by black dots with 0 size. PAS<1 (red) signifies the under-representation of the genes in the pathway in the given cell type, whereas PAS > 1 (violet) signifies the over-representation of the pathway. The mesophyll cells, which form multiple clusters, are represented by a single column in both panels. Pathways represented by a red dot are significantly under-represented in at least one cluster; pathways represented by a green dot are significantly under-represented in at least one cluster, and also significantly over-represented in at least one cluster. The size of the dots in the mesophyll column does not correspond to the *p*-value of PAS.

**Supplementary Fig. 10:**
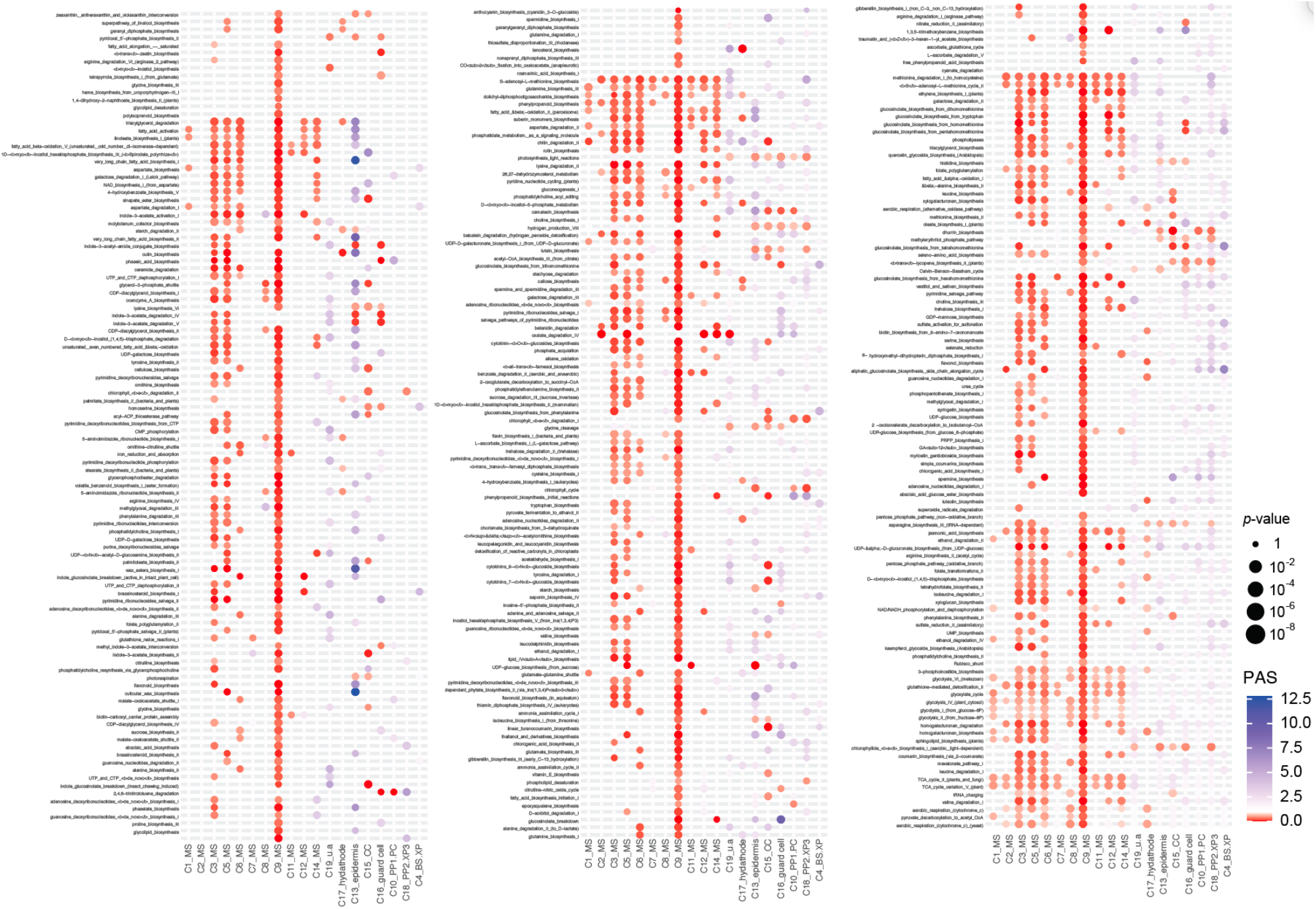
Metabolic pathway activity across all cell types. Statistical significance is represented as difference in dot size. Statistically insignificant values are left as blank (random permutation test, *p* >0.05). Colors represent pathway activity score (PAS); a score <1 (red) reflects a lower than average activity of the pathway in the given cell type, a score >1 (violet) indicates a higher activity.

**Supplementary Fig. 11:**
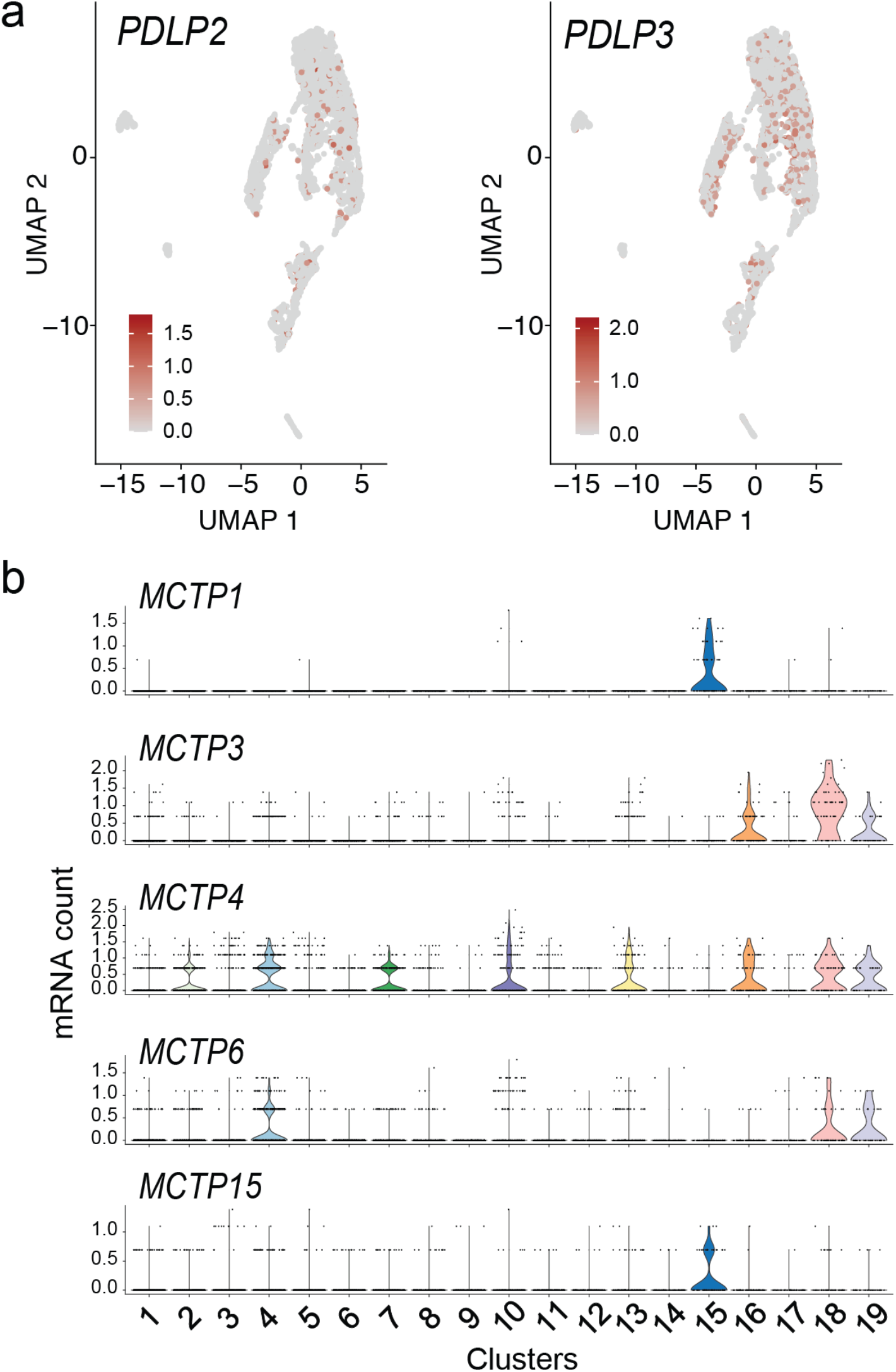
Cell type-specific transcript enrichment of plasmodesmatal proteins. **a**, UMAP plot showing the distribution of *PDLP2* and *PDLP3* transcripts. **b**, Violin plot showing transcript enrichment of *MCTP*s. Note that *MCTP1* and *FTIPI* (in Fig. 3) correspond to the same gene, AT5G06850.

**Supplementary Fig. 12:**
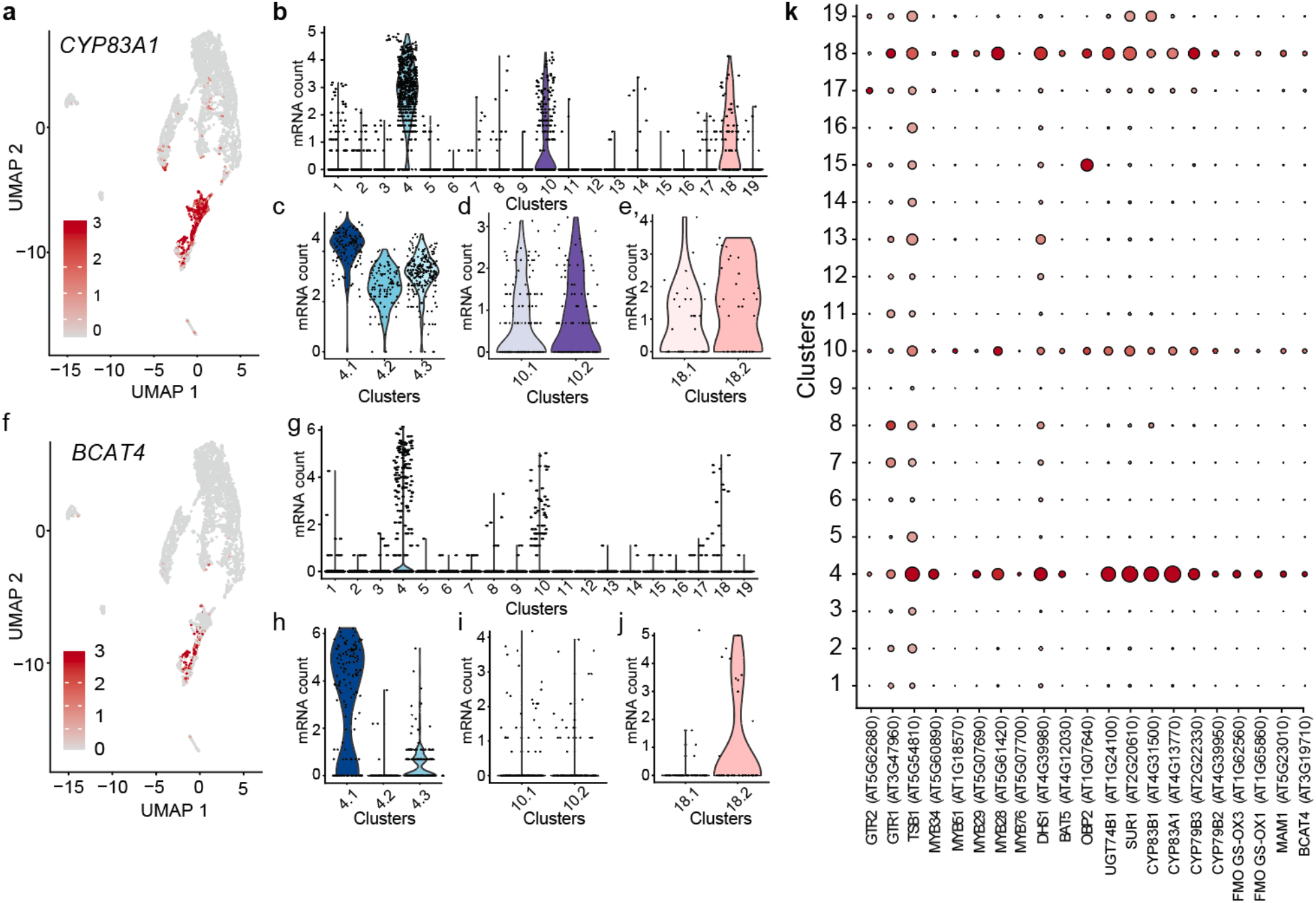
Enrichment of transcripts related to glucosinolate biosynthesis and transport. **a**, UMAP showing the enrichment of *CYP83A1* transcript. **b-e**, Violin plots illustrating the transcript enrichment of *CYP83A1* in BS1, BS2, XP **(b**,**c)**, PP1, PC^PP^, PC^XP^ **(b**,**d)**, PP2 and XP3 **(b**,**e). f**, UMAP showing the enrichment of *BCAT4* transcript. **g-j**, *BCAT4* transcript enriched in XP1, XP2 **(g**,**h)**, PP1, PC^PP^, PC^XP^ **(i)**, and XP3 **(j). k**, Dot plot showing enrichment of glucosinolate-related transcripts enriched in C4, C10, and C18. The diameter of the dot indicates the percentage of cells within a class, while the color encodes average enrichment across all cells within a class. These glucosinolate-related transcripts are known to be enriched in the BS, according to the translatome dataset^85^.

**Supplementary Fig. 13:**
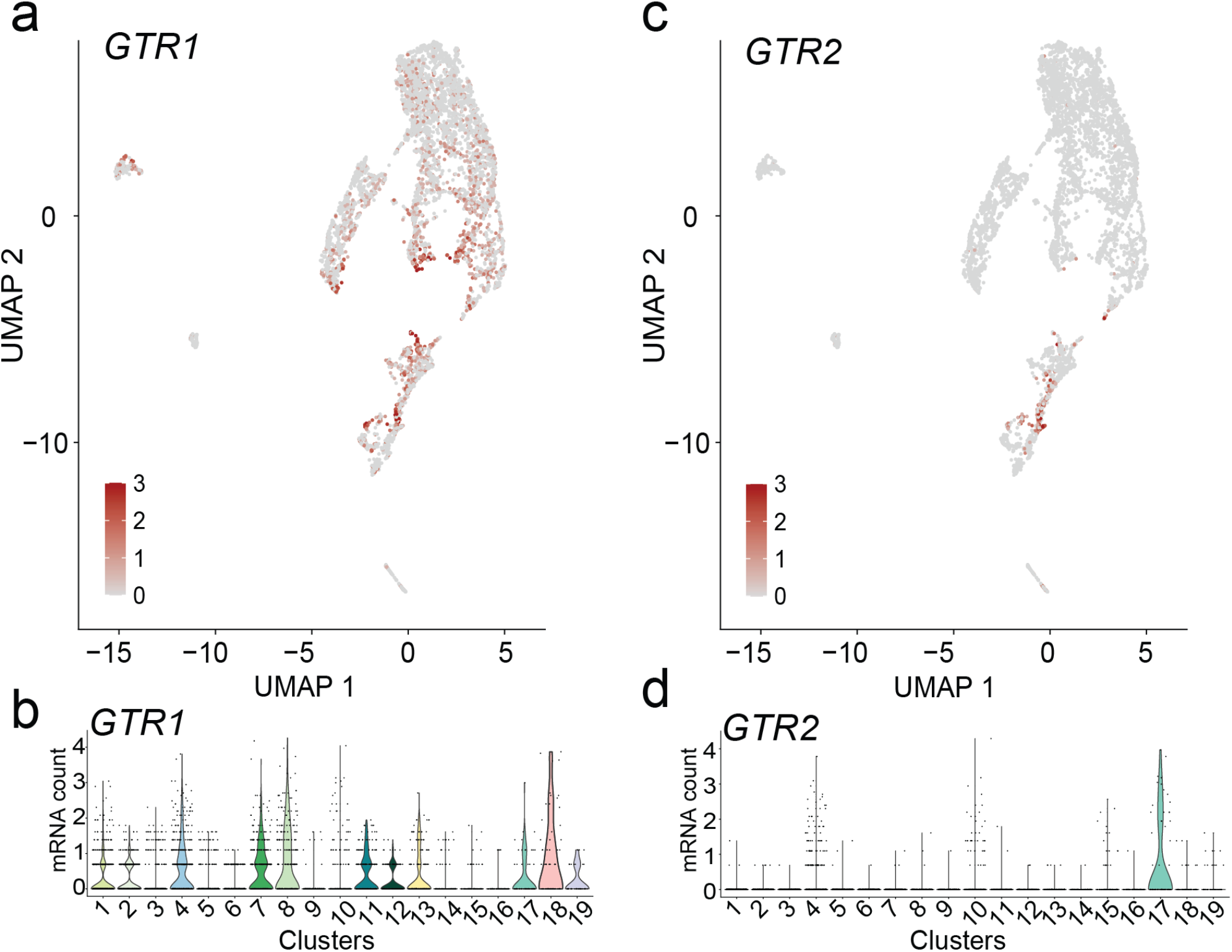
Enrichment of transcripts related to glucosinolate transport. **a**,**b**, *GTR1* transcript detected in the broadly across C4, C7, C8, C10, C11, C12, C13, C17, C18 and C19. **c**,**d**, *GTR2* transcript enriched in C4, C10, and a putative hydathode cluster, C17. *GTR1* and *GTR2* transcripts were not detected in phloem related clusters (CC, PP1, PP2) (**a**,**c**). The broad expression pattern of *GTR1* compared to *GTR2* has been previously reported^30^.

**Supplementary Fig. 14:**
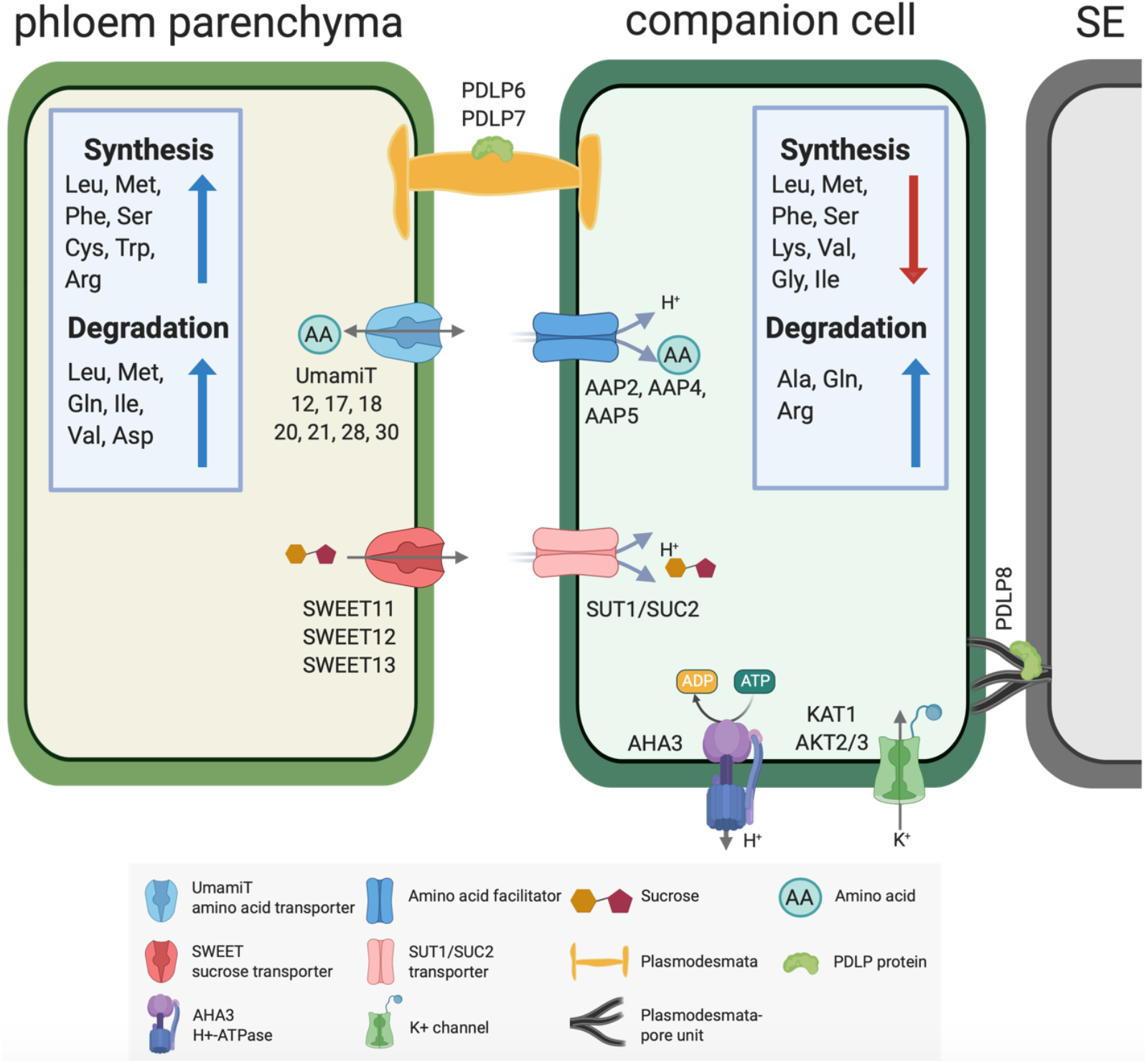
Hypothetical phloem loading process in Arabidopsis leaf. Sucrose produced during photosynthesis in the mesophyll cells is transported across the bundle sheath to the phloem parenchyma. Biosynthesis and catabolism of multiple amino acids is highly active in the phloem parenchyma. Transporters present in the phloem parenchyma secrete sucrose (SWEET11,12,13) or amino acids (UmamiT11, 12, 17, 18, 20, 21, 28, 30) into the apoplasm. H^+^/sucrose cotransporters import sucrose (SUT1/SUC2) and amino acids (AAP2,4,5) into the SE/CC. The H^+^ gradient required for the active import of sucrose and amino acids into the SE/CC is provided by plasma membrane H^+^-ATPases. The membrane potential is maintained by the potassium channels (KAT1 and AKT2/3). Symplasmic transport is mediated by PDLP6 and PDLP7 in the plasmodesmata in PP cells and PDLP8 enriched in the plasmodesmata-pore unit of the CC. Note that the schematic is based on transcript levels and the distribution could differ at the protein level. Schematics was made in ©BioRender – https://biorender.com

